# A Massively Parallel CRISPR-Based Screening Platform for Modifiers of Neuronal Activity

**DOI:** 10.1101/2024.02.28.582546

**Authors:** Steven C. Boggess, Vaidehi Gandhi, Ming-Chi Tsai, Emily Marzette, Noam Teyssier, Joanna Yu-Ying Chou, Xiaoyu Hu, Amber Cramer, Lin Yadanar, Kunal Shroff, Claire G Jeong, Celine Eidenschenk, Jesse E. Hanson, Ruilin Tian, Martin Kampmann

## Abstract

Understanding the complex interplay between gene expression and neuronal activity is crucial for unraveling the molecular mechanisms underlying cognitive function and neurological disorders. Here, we developed pooled screens for neuronal activity, using CRISPR interference (CRISPRi) and the fluorescent calcium integrator CaMPARI2. Using this screening method, we evaluated 1343 genes for their effect on excitability in human iPSC-derived neurons, revealing potential links to neurodegenerative and neurodevelopmental disorders. These genes include known regulators of neuronal excitability, such as TARPs and ion channels, as well as genes associated with autism spectrum disorder and Alzheimer’s disease not previously described to affect neuronal excitability. This CRISPRi-based screening platform offers a versatile tool to uncover molecular mechanisms controlling neuronal activity in health and disease.

## Introduction

Billions of neurons in the human brain form a complex, dynamic network that gives rise to the emergent properties of cognition, memory, and behavior. Different neuronal cell types in the brain are defined by distinct patterns of gene expression,^1–4^ with recent studies demonstrating these genetically defined subtypes can have distinguishable spike patterns.^5^ Neuronal activity, in turn, further alters gene expression to affect the composition of synaptic receptors and ion channels at the neuronal membrane, making activity-dependent gene expression critical for long-lasting plasticity.^6,7^ Large-scale efforts to map gene expression to different areas of the brain,^8,5^ at single cell resolution^9,10^, or to connect transcriptional changes to electrophysiological measurements^11^ are underway, but we lack a systematic understanding of how expression differences of individual genes affect neuronal activity.

One approach to link genes to neuronal activity and brain function is to map genetic risk factors underlying neurological disorders that are characterized by neuronal activity changes. In epilepsy, human genetics have revealed mutations in voltage-gated ion channels (channelopathies), synaptic channels (synaptopathies), and metabolic genes in both sporadic^12^ and familial^13^ disease. Other diseases, such as Alzheimer’s Disease (AD) and autism spectrum disorder, also involve changes in neuronal activity. However, it is unclear which of the associated risk genes^14,15^ affect neuronal activity, and we lack a systematic and scalable approach to establish the effect of genes on neuronal function.

Classically, interrogation of neuronal activity mechanisms has been accomplished through patch-clamp electrophysiology. This technique is the gold-standard to measure a range of properties, including spontaneous synaptic currents and individual channel kinetics. While powerful, these techniques are limited by the number of neurons that can be interrogated as the researcher must make physical contact with each individual neuron. This requires both a high degree of technical skill and large time investment to evaluate each perturbation.

To measure electrical activity from dozens of neurons in a network simultaneously, researchers have turned to the use of multi-electrode arrays (MEAs) and fluorescent activity probes. MEAs circumvent some of the drawbacks of patch-clamp electrophysiology by measuring local field potentials in a network of neurons but are costly and often sacrifice single-neuron spatial resolution. Alternatively, fluorescence-based indicators of calcium^16^ and membrane potential^17,18^ also circumvent the throughput constraints of patch-clamp electrophysiology while still providing single-neuron resolution. Furthermore, optogenetics using microbial-opsins provides additional temporal control and the ability to stimulate specific neurons.^19,20^ Still, the number of perturbations, genetic or pharmacologic, that can be performed are limited in that each must be done in an arrayed format on high-content imaging plates. Recently, it was shown that calcium oscillations detected using GCaMP6 could be used for multiparametric drug screening in healthy and SOD1^A4V^ motor-neurons differentiated from human iPSCs,^21^ however, to our knowledge, a method capable of comprehensively evaluate genetic modifiers of activity on a genome-wide scale has not been reported.

To enable the scalable characterization of gene function in human cells, we developed CRISPR interference (CRISPRi) and activation (CRISPRa) screening^22^ and implemented it in human iPSC-derived cell types relevant for brain function and diseases, including neurons,^23,24^ microglia,^25^, astrocytes^26^, and neuron-glia iAssembloids^27^. These screens allow for unbiased identification of genes and mechanisms underlying cell type specific biological processes. While pooled CRISPR-based screens have been successfully deployed to elucidate neuronal phenotypes such as survival,^23,24,27^ transcriptomic states,^23,24^ oxidative damage,^24^ lysosomal function,^24^ and protein aggregation,^28^ it has so far not been possible to use pooled CRISPR-based screens to uncover modifiers of neuronal activity. Here, we report a method to uncover modifiers of neuronal excitability in massively parallel screens.

To overcome the challenges in evaluating modifiers of neuronal activity at scale, we developed a pooled screening approach using integration of calcium signal. The fluorescent calcium integrator CaMPARI2 acts as an irreversible record of calcium levels in a neuron by photoconverting while in the presence of high calcium and UV light.^29–32^ Based on the ratio of photoconversion in each neuron, the relative level of activity can be discerned by flow cytometry. CaMPARI2 was previously used in a genetic screen to identify modifiers of calcium entry in mouse embryonic fibroblasts^33^. Here, we CaMPARI2 in human iPSC-derived neurons in conjunction with CRISPRi to conduct pooled genetic screens and uncover modifiers of neuronal excitability. We validated hit genes using calcium imaging and patch-clamp methods and generated hypotheses for how these hit genes regulate neuronal excitability.

## Results

### CaMPARI2 photoconversion provides a ratiometric readout for glutamate-induced activity in human iPSC-derived neurons

To establish a readout of neuronal activity compatible with pooled CRISPR-based screens using fluorescence-activated cell sorting (FACS), we evaluated the utility of the fluorescent calcium integrator CaMPARI2.^32^ We introduced a CaMPARI2 transgene into our iPSC line expressing CRISPRi machinery and inducible Ngn2^23^ using lentiviral integration followed by clonal selection (Figure 1a). These iPSCs are rapidly differentiated into glutamatergic neurons by doxycycline-induced expression of NGN2, and express dCas9-KRAB for efficient knockdown of genes targeted by a sgRNA.^23^

**Figure 1.**
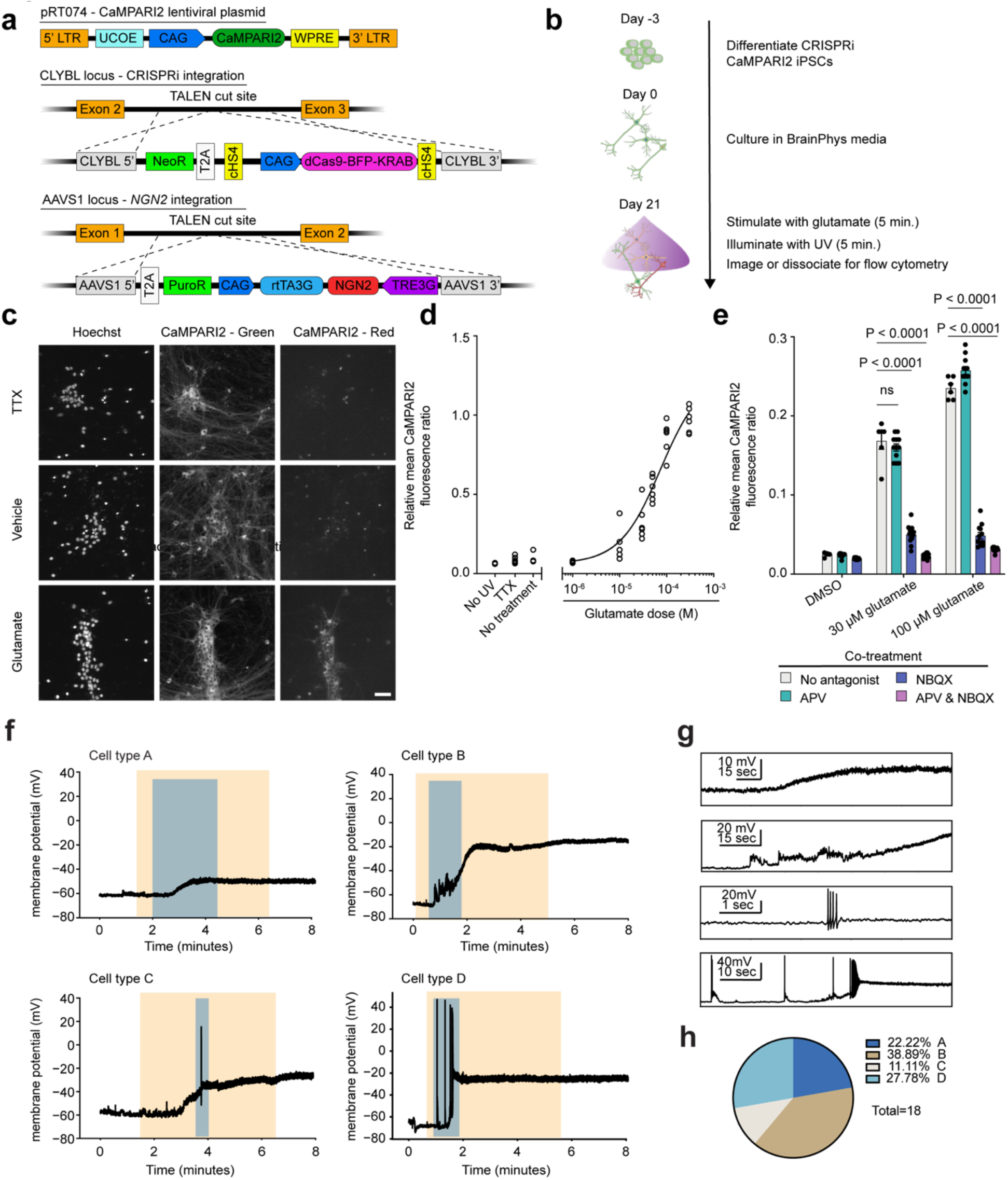
CaMPARI2 captures glutamate receptor activation in iPSC-derived glutamatergic neurons. (a) Strategy for generating CRISPRi-NGN2-CaMPARI2 iPSC line: NGN2 and dCAS9-KRAB was stably integrated into the AAVS1 and CLYBL loci, respectively, as described previously.17 CAG promoter-driven CaMPARI2 was randomly inserted into the genome via lentiviral integration. UCOE: Ubiquitous Chromatin Opening Element, WPRE: Woodchuck Hepatitis Virus Posttranscriptional Regulatory Element. (b) Experimental workflow of CaMPARI2 photoconversion assay in glutamatergic NGN2 neurons. (c) Representative fluorescence micrographs of iPSC-neurons expressing CaMPARI2 following incubation with 1.5 µM tetrodotoxin (TTX), vehicle (0.1% DMSO), or 30 µM glutamate for 5 minutes and illumination with 405 nm light. Nuclei were stained with Hoechst 33342. Unconverted CaMPARI2 persists in the soma and neurites, and converted CaMPARI2 is only observed with high signal after incubation with 30 µM glutamate. Merged micrographs show colocalization of nuclei (blue), CaMPARI2 (green), and converted CaMPARI2 (magenta). Scale bar is 20 µm. (d) CaMPARI2 neurons respond to glutamate in a dose-dependent manner. N = 6 independent culture wells per measurement. “No UV”, TTX (1.5 µM), and “no treatment” conditions did not contain glutamate. (e) Co-incubation of glutamate with AMPA receptor (10 µM NBQX), but not NMDA receptor (50 µM APV), antagonists attenuate CaMPARI2 photoconversion, suggesting that AMPA receptor activation is the primary mechanism of calcium influx. N = 6 independent culture wells per measurement. Unpaired t-test, * = P <0.0001. Error bars are SEM. (f) Representative current clamp recordings from four different cell types. The orange shaded area indicates the 5-minute window when 30 µM glutamate was applied to the recording chamber. The blue shaded area highlights the zoomed-in view shown in (g) for each trace. (g) From top to bottom, a zoomed-in view of the blue shaded areas from (f) for Cell types A to D. (h) Percentage of each cell type from recorded iNeurons. N = 18 neurons.

Synaptic dysfunction is a critical hallmark of many human neurological diseases,^34–38^ therefore we focused our efforts on establishing a CaMPARI2-readout for synaptic excitability. To depolarize glutamatergic synapses, we stimulated iPSC-neurons with glutamate after 21 days of differentiation and illuminated with UV light to photoconvert CaMPARI2. We then measured the CaMPARI2 red-to-green fluorescence ratio after glutamate stimulation using both microscopy and flow cytometry (Figure 1b). With confocal microscopy, we found a robust increase in CaMPARI2 photoconversion in neurons treated with glutamate compared to vehicle control (Figure 1c). Virtually no photoconversion was observed with glutamate treatment in tetrodotoxin (TTX)-treated neurons (Figure 1c). TTX-treated neurons showed similar signal compared to vehicle control suggesting that the baseline synaptic excitability of iPSC-neuron cultures is low.

We tested the dose-response relationship of glutamate and CaMPARI2 photoconversion by flow cytometry (Figure 1d) and found that incubation with 30 µM glutamate for 5 minutes followed by exposure to UV light provided good signal without reaching saturation. This intermediate concentration was chosen to provide dynamic range to enable identification of genetic modifiers that either decrease or increase excitability. Co-incubation of glutamate with APV or NBQX, antagonists for NMDA and AMPA receptors, respectively, demonstrated that the detected calcium release is primarily through AMPA receptors (Figure 1e). Additionally, co-incubation of neurons with glutamate and metabotropic glutamate receptor (mGluR) antagonists modulated neuronal excitability, but full inhibition of signal was achieved only in the presence of APV and NBQX (Extended Data Figure 1b). Treatment with KCl also induced a large CaMPARI2 signal (Extended Data Figure 1c-d), demonstrating that multiple methods of neuronal activation can be interrogated with CaMPARI2. Furthermore, we found that 5 minutes of UV illumination maximized signal (Extended Data Figure 1e). Neuron plating density had a slight effect on the photoconversion outcome at 21 days of differentiation (Extended Data Figure 1f). Five minutes of UV illumination had minimal effects on cell viability during the experimental timeline (Extended Data Figure 1g-h) and treatment with glutamate did not cause acute toxicity (Extended Data Figure 1i).

To better understand how our iNeuron system responds to glutamate, we performed current-clamp recordings while treating with 30 µM glutamate for 5 minutes. We find that iNeurons have a heterogeneous response to glutamate, which we classify into four distinct voltage response profiles (Figure 1f). These include iNeurons that do not fire action potentials when treated with glutamate, including those with small depolarizations (Cell type A; N = 4, 22.2% of total cells), and those with larger depolarizations to membrane potentials above –40 mV (Cell type B; N = 7, 38.9% of total cells). Additionally, we found iNeurons that depolarize and fire action potentials, and upon glutamate wash-out repolarize (Cell type C; N = 2, 11.11% of total cells) or remain depolarized (Cell type D; N = 5, 27.8% of total cells).

Current injection data from these iNeurons before glutamate stimulation show the heterogeneity of intrinsic membrane properties that correlate with the different types of responses to glutamate stimulation (Extended Data Figure 2a-b). Cell types A and B showed a trend towards higher rheobase compared to Cell types C and D (Extended Data Figure 2c, one-way ANOVA, p = 0.05). Although Cell types A and B did not generate action potentials with glutamate stimulation, they were capable of firing action potentials with high amplitude current injections. Overall, we found that 30 µM glutamate stimulation for 5 minutes generates diverse responses in iNeurons.

### CRISPRi screen for modifiers of neuronal excitability

With this technology in hand, we focused our efforts to study neuronal excitability, which has been implicated in many neurological disorders. Patients with Alzheimer’s Disease (AD) have an 8-to 10-fold increase in the risk of seizures,^39,40^ with familial early-onset cases having more severe risk,^39^ and seizure prevalence has been associated with accelerated cognitive decline.^41^ Furthermore, neuronal network dysfunction has been proposed as a critical contributor to AD that occurs decades before clinical symptoms manifest.^42^ Epilepsy also shows comorbidity with 5-46% of individuals with autism spectrum disorder (ASD),^38^ which is believed to be caused by an imbalance of synaptic excitation and inhibition (E/I balance) or dysfunctional ion channels. Identifying the genetic etiology can help inform precise treatment options for patients;^13,43^ however, many of these cases are resistant to current anti-seizure medications and monogenic syndromes are only identified in about 40% of patients with epilepsy.^13^ This highlights the need for methods that comprehensively evaluate the functional consequences of genetics on neuronal activity.

To establish CRISPRi screening based on the CaMPARI2 reporter, we generated a custom sgRNA library targeting 1,343 genes selected based on their known or proposed roles in neuronal activity (Supplementary Data Table 1), neurodegeneration, and neurodevelopmental disorders (Supplementary Data Table 2). We designed this library to evaluate our screening paradigm by knocking down known regulators of neuronal excitability and potentially uncover new roles for genes implicated in human diseases. The library targets each gene with 5 independent sgRNAs and contains 250 non-targeting control sgRNAs, resulting in a library of 7,450 different sgRNAs (Supplementary Data Table 3).

We transduced our CaMPARI2-CRISPRi-Ngn2 iPSCs with the lentiviral sgRNA library, selected for transduced iPSCs using puromycin, expanded, and then differentiated into neurons by inducing NGN2 expression with doxycycline (Figure 2a). After 21 days of differentiation, we stimulated neuronal activity with 30 µM glutamate, photoconverted CaMPARI2 with UV light for five minutes using a UV light box, and then dissociated neuronal cultures for FACS. We collected neurons with the highest 35% and lowest 35% CaMPARI2 red/green fluorescence ratio, justified previously by extensive quantitative modeling.^44^ We determined the frequency of neurons expressing each sgRNA in the library using next-generation sequencing to determine which sgRNAs cause significant increase or decrease in CaMPARI2 response to glutamate exposure (Figure 2a). We performed this screen in duplicate to identify the most robust hits.

**Figure 2.**
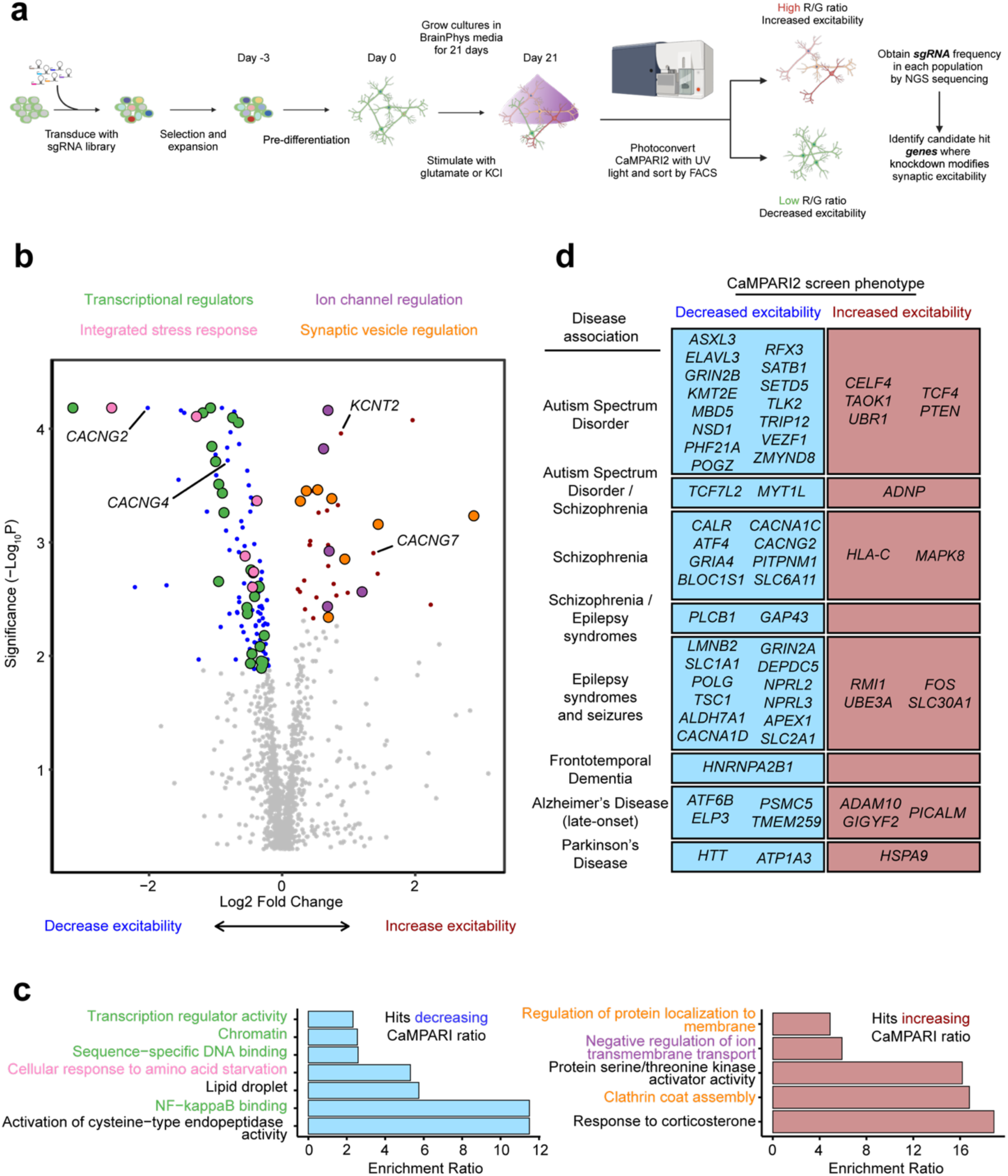
CRISPR interference (CRISPRi) screening with CaMPARI2 reveals genetic modifiers of neuronal excitability in iPSC derived neurons. (a) Screening strategy using CaMPARI2 as readout for neuronal excitability. CRISPRi iPSC expressing CaMPARI2 were transduced with a lentiviral sgRNA library focused on genes annotated for neuronal activity, neurodegenerative disease, or neurodevelopmental disorders (7449 sgRNA constructs targeting 1343 genes). Exposure to puromycin selects for iPSCs successfully transduced with sgRNA construct. iPSCs were then differentiated into neurons via overexpression of NGN2 and cultured for 21 days. Neurons were then incubated with 30 µM glutamate for five minutes, illuminated with UV light for five minutes, and then separated by FACS into high and low red/green fluorescence ratio populations. Frequency of neurons expressing sgRNAs were determined by next-generation sequencing for high and low red/green ratio neuronal populations and hit genes were identified. (b) Volcano plot summarizing the effect of knockdowns on CaMPARI phenotypes and the determine statistical significance for targeted genes (Mann-Whitney U test). Dashed lines indicate a false-discovery rate (FDR) cutoff of 0.05, based on the phenotype score for genes calculated from 5 targeting sgRNAs. Red and blue points indicate hit genes where knockdown increases and decreases CaMPARI2 red/green fluorescence ratio, respectively, in response to glutamate stimulation. Grey points indicate non-hits. Large colored dots indicate hit genes that were found in the enriched GO terms in (d). (c) Disease-associated genes that influenced excitability in the CaMPARI2 screen. (d) Over-representation analysis (ORA) using Webgestalt indicates gene ontology (GO) annotation terms that describe genes enriched in the low (blue) and high (red) CaMPARI2 ratio from the screening library.

Of the 1,343 genes targeted by our custom sgRNA library, we identified 116 genes for which knockdown decreased synaptic excitability and 37 genes for which knockdown increased synaptic excitability, for a total of 153 hits (Figure 2b, FDR 0.1). We found that the observed hits have Gene Ontology (GO) annotations related to neurotransmitter receptor activity, ion transport, and synaptic transmission (Extended Data Figure 3a).

We performed an Over-Representation Analysis (ORA) using WebGestalt^45^ to identify enrichment of functional classes of genes among the hits (Figure 2b,c). Of the hits that increased synaptic excitability, GO terms for membrane protein localization, clathrin coat assembly, and negative regulation on ion transmembrane transport were over-represented, potentially highlighting mechanisms related to neuronal homeostasis at the plasma membrane and synaptic vesicle recycling. Additionally, many GO terms related to gene transcription and chromatin regulation (Figure 2c) were over-represented in hits that decrease synaptic excitability, which suggests altered gene expression is important for synaptic function or could be related to delayed neuronal differentiation. The GO term *Cellular Response to Amino Acid Starvation*, an arm of the integrated stress pathway (ISR), was over-represented in hits decreasing synaptic excitability. Previous studies have explored the use of ISRIB, and ISR inhibitor, to restore cognitive deficits in mice.^46^ Furthermore, our screen revealed that knockdown of *EIF2AK3, EIF2AK4,* and *ATF4* reduced excitability, in line with previous work showing the role of eIF2a in long-term memory formation.^47,48^ These hit genes highlight that neuronal excitability is influenced by diverse molecular pathways and that the CaMPARI2-based screening method is capable of detecting neuronal excitability modifiers that are upstream in these pathways.

Many hit genes have been associated with Alzheimer’s Disease, epilepsy, and/or autism spectrum disorder (Figure 2d, Extended Data Figure 3b). While our custom library was purposely enriched with these disease-associated genes (Supplementary Data Table 2), this screen suggests a potential, previously unknown role these hit genes play in regulating neuronal excitability.

### Validation of screen hits reveals mechanisms of altered excitability including altered synaptic function and action potential dynamics

We examined these observed screen phenotypes by flow cytometry using two individual guides targeting *CACNG2* and *KCNT2*, directly comparing the CaMPARI2 ratio after glutamate stimulation of iPSC-neurons receiving the sgRNA versus iPSC-neurons without guide (Figure 3a). This is expressed as a normalized CaMPARI ratio (neuronal excitability), where the mean response of neurons in wells with no sgRNA transduction is equal to one. We evaluated neuronal networks in which 100% of neurons received a sgRNA and networks in which 50% of neurons received a sgRNA which allowed us to evaluate both cell autonomous and cell non-autonomous effects on neuronal excitability.

**Figure 3.**
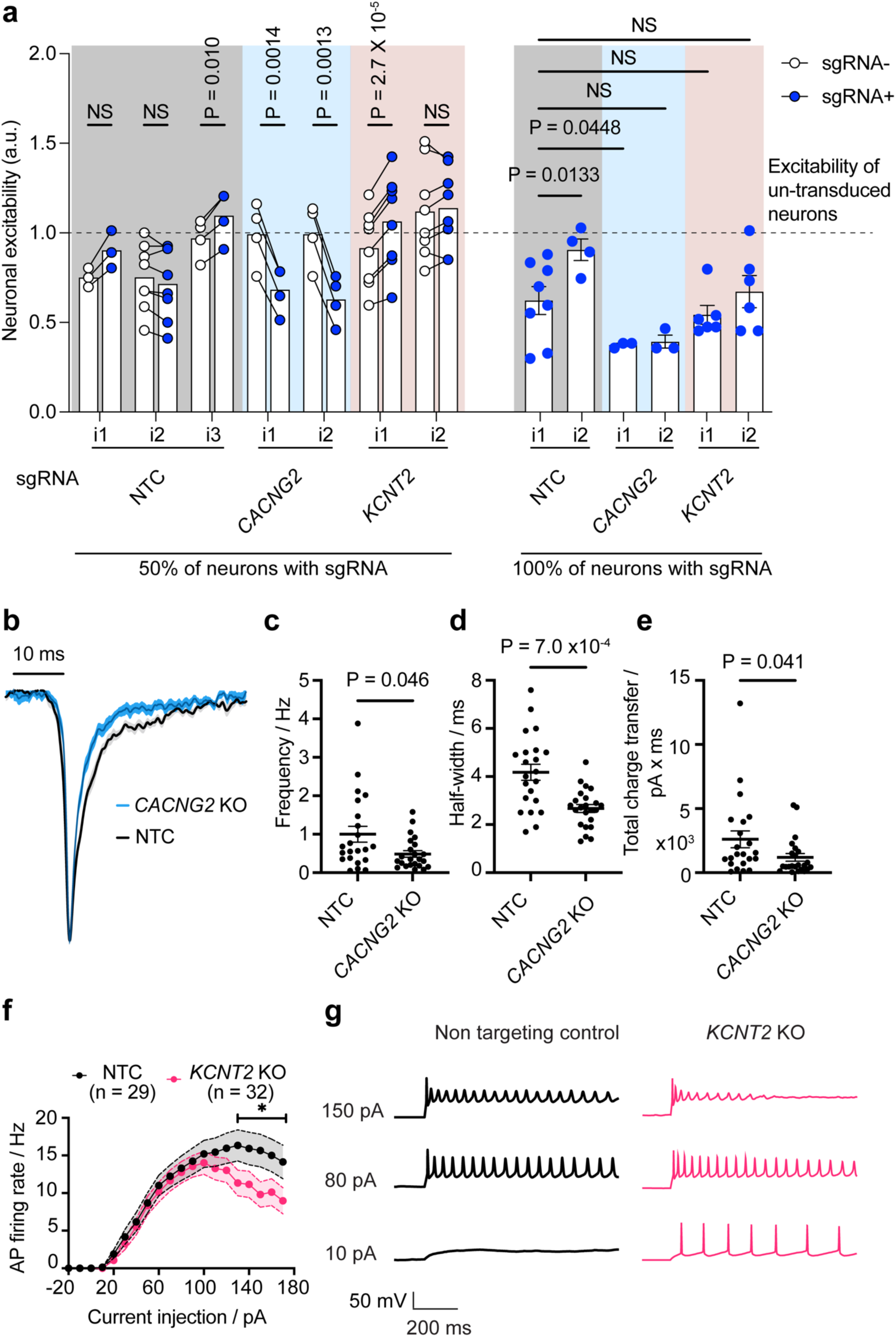
CACNG2 and KCNT2 modify excitability in human neurons. (a) CaMPARI2 neurons expressing sgRNAs targeting CACNG2 have reduced neuronal excitability in response to glutamate stimulation. This effect is observed intra-well, where CACNG2 sgRNA-containing neurons have a lower CaMPARI2 ratio than those that do not receive the sgRNA, and inter-well, in cultures in which 100% of the neurons are expressing the sgRNA. One of the sgRNAs targeting KCNT2 shows an increase in neuronal excitability when comparing intra-well effects, which matches the screening phenotype. In cultures where 100% of the neurons are expressing one of the sgRNAs targeting KCNT2, there is an unexpected decrease in the observed neuronal excitability. CaMPARI2 ratios are normalized to neuron samples that do not receive a sgRNA and are run on the flow cytometer on the same day. 50% knockdown cultures: N = 3-8 independent culture wells per experiment. Each individual experiment was normalized to the excitability (CaMPARI2 response) of wells where neurons did not receive a sgRNA. For intra-well comparisons, a paired ratio t-test between sgRNA– and sgRNA+ populations were performed for each sgRNA. The calculated P values are corrected for multiple hypothesis (Holm-Šídák method). For inter-well comparisons, P values were calculated by ANOVA, using NTC_i1 as reference. P > 0.05 = not significant (NS). (b) The spontaneous excitatory postsynaptic currents (sEPSCs) in CACNG2 KO iNeurons demonstrate accelerated kinetics and a decrease in overall charge transfer. Averaged and normalized sEPSCs from all the recorded instances of NTCs (N = 22, in black) and CACNG2 KOs (N = 23, in blue), with the shaded area indicating standard errors. (c) CACNG2 KOs exhibit reduced sEPSC frequency, (d) half-width, and (e) total charge transfer (Mann-Whitney test). NTCs (N = 22) and CACNG2 KOs (N = 23). Error bars denote standard error. (f) KCNT2 KO neurons exhibit premature action potential failure. Current-spike output relationship presenting a diminished action potential firing rate in KCNT2 KOs (N = 32) at high current injection amplitudes, as juxtaposed against non-targeting controls (N = 29). The differences were statistically significant for current injection steps greater than 120 pA (asterisk donates p values as follows: 130 pA: p = 0.01, 140 pA: p = 0.02, 150 pA: p <0.01, 160 pA: p = 0.02, 170 pA: p = 0.01; two-way ANOVA with multiple comparisons). (g) Sample recordings from both NTCs and KCNT2 KOs illustrating voltage responses of neurons to varying degrees of current injection.

Knockdown of *CACNG2* decreased neuronal excitability in response to glutamate when compared to neurons that were not transduced with the sgRNA (Figure 3a). We tested the effect of this knockdown in neuronal populations where approximately 50% of the neurons expressed an sgRNA and found a robust decrease in excitability in *CACNG2* knockdown neurons compared to intra-well neurons without a sgRNA. In populations where approximately 100% of the neurons had *CACNG2* knockdown, neuronal excitability was dramatically reduced when compared to populations of neurons without sgRNA treatment (Figure 3a). Similarly, with *KCNT2* knockdown we observed increased excitability as observed in the screen (Figure 3a). In populations where all neurons were transduced, there was an overall decrease in excitability, which opposes the screen phenotype and suggests more complicated roles for *KCNT2* via both cell autonomous and network level impacts. Together, these results suggest that the CaMPARI2 screening modality can capture genetic modifiers of activity.

Our CaMPARI2-based CRISPRi screening method positively identified modifiers of neuronal excitability, including AMPAR (*CACNG2, GRIA4*) and NMDAR (*GRIN2A*, *GRIN2B*) subunits and ion channels (*KCNT2, CACNA1C, CACNA1D*). To explore neuronal excitability mechanisms that could be detected by the CaMPARI2 screening approach, we evaluated the effects of *CACNG2* and *KCNT2* perturbation using whole-cell electrophysiology. For these experiments, ribonucleoprotein (RNP) mediated CRISPR was used to generate knockouts (KOs) in human iPSC neurons, with NTC guides as negative controls. Recordings of spontaneous excitatory postsynaptic currents (sEPSCs) revealed differences in synaptic activation between *CACNG2* KO neurons and neurons treated with NTCs (Figure 3b, Extended Data Figure 4c,d). A small reduction in the frequency of detected sEPSCs in *CACNG2* KOs was observed compared to NTC (Figure 3c). Furthermore, the half-width and charge transfer of these sEPSCs was significantly attenuated in *CACNG2* KO neurons (Figure 3d-e), reflecting altered receptor kinetics. This decreased synaptic activation observed in the whole cell recordings aligns with previous observations,^49–54^ and reflects the reduced excitability observed in the CaMPARI2 screen. These results indicate that the CaMPARI2 screening approach is sensitive to detect known regulators of synaptic excitability with complex electrophysiological phenotypes.

We observed no effect on passive membrane properties in *KCNT2* KO neurons (Extended Data Figure 4g-i). However, large current injections caused a drop in action potential firing rate in *KCNT2* KO neurons and premature entry into depolarization block, whereas neurons with a NTC sgRNA do not (Figure 3f-g). These results indicate that in addition to synaptic regulators, the CaMPARI2 screening approach is sensitive to detect regulators of action potential firing such as ion channels.

### Modifiers identified in the CaMPARI2 screen are reproducible, specific to neurons, and can be stimulus-specific

We generated a focused CRISPRi sgRNA library targeting 395 hit genes (Supplementary Table 4) from our primary screen (based on a relaxed criterion for hit genes, see Methods) for secondary validation screens. We subjected CaMPARI2 neurons to either treatment with 30 µM glutamate or 50 mM KCl in two independent screens (Extended Data Figure 3a). We selected 50 mM KCl treatment as it resulted in a CaMPARI2 response with large dynamic range (Extended Data Figure 1c-d). Comparison of gene scores from our primary glutamate-based screen (Figure 2) and our secondary glutamate-based validation screen showed a high degree of correlation (Pearson’s R = 0.60, Figure 4a), suggesting that the CAMPARI2 screening paradigm produces reproducible results. Many genes had similar effects when knocked down in the secondary glutamate-stimulated screen and the KCl-stimulated screen (Pearson’s R = 0.33, Figure 4b), suggesting mechanisms that modify neuronal excitability in a general way irrespective of the depolarization method. Still, there were many hits that specifically modified excitability in the secondary glutamate screen, but not in the KCl screen (Extended Data Figure 6f). For instance, *KCNT2* knockdown had no significant phenotype with KCl-mediated depolarization, despite our observation that *KCNT2* KO neurons enter depolarization block prematurely during high levels of current injection (Figure 3f-g).

**Figure 4.**
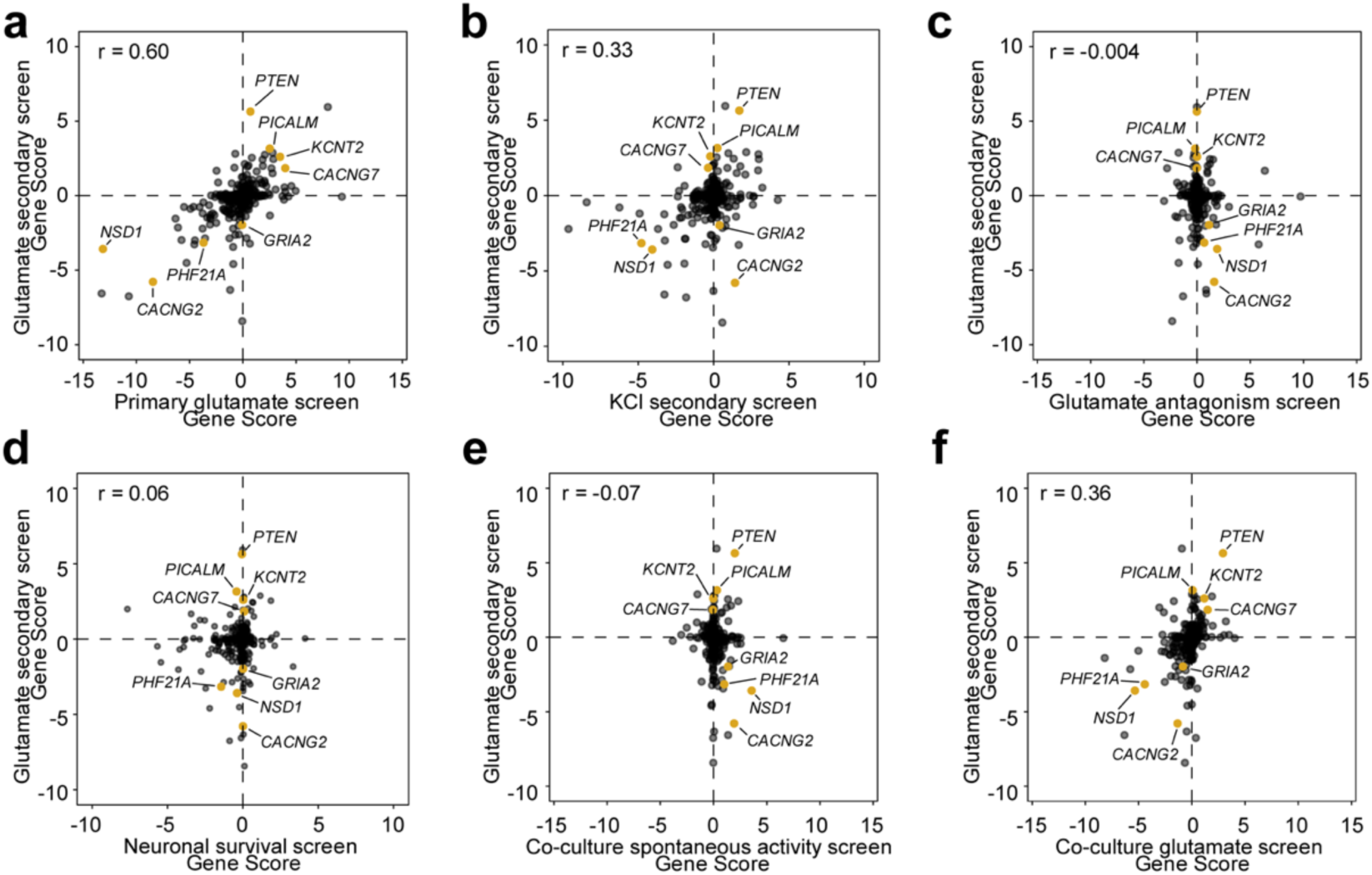
Secondary CRISPRi screens reveal excitability phenotypes in iPSC neurons. (a) Comparison of gene scores across different screens. Gene scores of knockdowns in the primary and secondary screens with glutamate stimulation show a positive correlation (Pearson’s R = 0.60). (b) Many hit genes show similar phenotypes in neurons depolarized with KCl compared to those stimulated with glutamate (Pearson’s R = 0.33). (c) Glutamate receptor antagonists mostly abrogate phenotypes in neurons stimulated with glutamate, as reflected by a weak correlation of gene scores (Pearson’s R = –0.004) (d) Excitability phenotypes in glutamate-stimulated neurons are not strongly correlated with survival phenotypes (Pearson’s R = 0.064). (e) Modifiers of activity of spontaneously firing neurons in coculture do not correlate with those of glutamate-stimulated neurons in monoculture (Pearson’s R = –0.07). (f) Gene scores of knockdowns in the secondary screens with glutamate stimulation in monoculture and coculture show a positive correlation (Pearson’s R = 0.36).

To evaluate if there are modifiers that primarily affect neuronal calcium homeostasis instead of activity, we co-treated CaMPARI2 neurons with 30 µM glutamate and an antagonist cocktail for full GluR blockage (Extended Data Figure 5b). While we found some modifiers that had a statistically significant gene score, comparison of gene scores to the secondary glutamate screen showed no correlation (Pearson’s R = –0.004). This would suggest that while some modifiers influence neuronal calcium homeostasis, the underlying signal from this effect is overshadowed by treatment with glutamate and the effects on neuronal excitability.

We collected neurons at the start of differentiation (day 0) and after 21 days of differentiation (before stimulation and photoconversion) to identify knockdowns that affect neuronal survival (Extended Data Figure 5c). To assess which hit genes are specific to glutamate-stimulated activity in neurons, we also treated a subset of the sgRNA-transduced undifferentiated iPSCs with glutamate and used the same photoconversion and FACS strategy (Extended Data Figure 5d). In the survival screen, we found 90 significant hit genes with a toxic knockdown phenotype in neurons, and 16 protective hits. Comparison to the secondary glutamate screen (Figure 4d) showed little correlation of these effects (Pearson’s R = 0.064). Furthermore, we did not identify any significant hits in iPSCs treated with glutamate. Together, these results indicate that the CaMPARI2 screen was specific in detecting modifiers of neuronal excitability.

While the use of monoculture iPSC-neurons provides a convenient, reductionist system for biological discovery, the physiological immaturity of these neurons make neuronal excitability changes difficult to relate to more advanced systems. It has been established that coculture of iPSC neurons with astrocytes improves the electrophysiological maturation of the neurons. Therefore, we evaluated changes to neuronal excitability in a coculture of the NGN2 neurons with human iPSC-astrocytes (iAstrocytes)^26^ by comparing CaMPARI2 signal by flow cytometry and found a slight increase in spontaneous activity signal and higher sensitivity to glutamate (EC_50_ = 12 µM; Extended Data Figure 6a). We also observed a higher spike count in cocultures relative to monocultured iPSC neurons grown on multi-electrode arrays (MEAs) as early as 14 days in culture (Extended Data Figure 6b). Furthermore, the synchrony and spike number of these cocultures is disrupted after treatment with tetanus toxin (TeNT; Extended Data Figure 6c-d), suggesting the presence of synaptic transmission.

To assess the ability of the CaMPARI2 system to evaluate modifiers of neuronal excitability in this coculture model, we examined the effect of these modifiers in spontaneously active cultures and when treated with glutamate (Extended Data Figure 5e). There was no correlation of modifiers affecting spontaneous activity in cocultured neurons when compared to the glutamate treated monocultured iPSC-neurons (Figure 4e). However, performing the screen in coculture and using glutamate has a positive correlation (Pearson’s R = 0.36) to the screen performed in monocultured neurons (Figure 4f). Knockdown of synaptic genes like *CACNG2* and *GRIA2* decreased excitability in response to glutamate in both monoculture and coculture, while the *KCNT2* KD increased excitability in both. We noticed genes with stronger phenotypes in monoculture than in coculture (i.e*. KCNJ9,* Extended Data Figure 6g) and cases where phenotypes were stronger in coculture (i.e. *GRIN2B*, Extended Data Figure 6g). This may reflect the changes in excitability between culture conditions, however additional mechanistic validation is needed.

### Evaluating mechanisms of disease-associated modifiers of excitability

We selected six additional hit genes that have been associated with human neurological disease for additional validation (Figure 5). *NSD1* and *PHF21A*, both of which decreased synaptic excitability in the glutamate screens, are histone modifiers that have been identified as risk genes for ASD. Comparing the CaMPARI2 response in neurons receiving sgRNAs normalized to neurons without a guide, individual knockdown of *NSD1* and *PHF21A* recapitulated the observed phenotype in the screen (Figure 5a). We also validated the screening phenotype for *GRIA2* (Figure 5a), an AMPA receptor subunit that is downregulated after kainate-induced status epilepticus in the hippocampus.^55,56^

**Figure 5.**
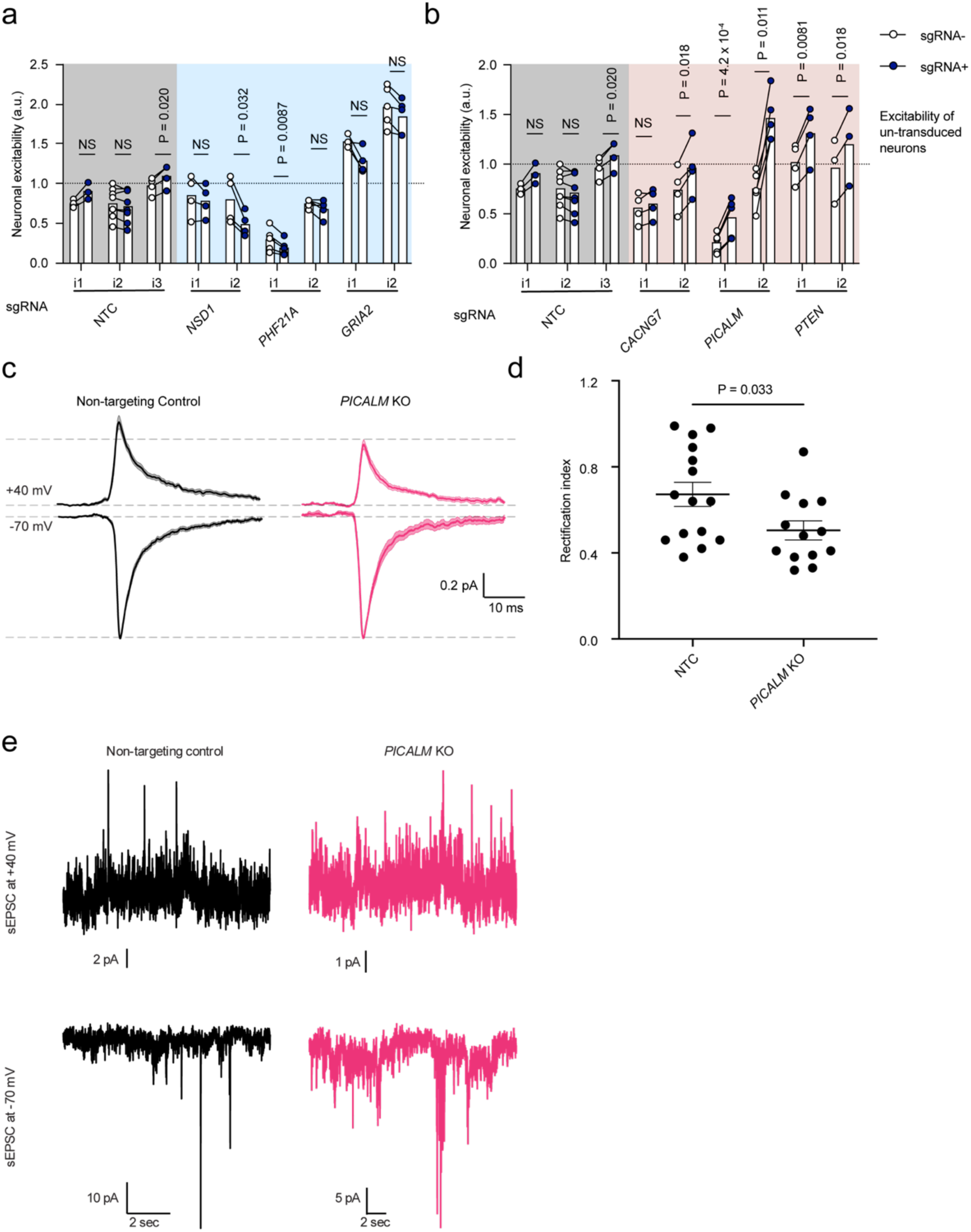
Flow validation of disease-associated excitability hit genes. (a) CaMPARI2 neurons expressing sgRNAs targeting genes that, when knocked down, reduced neuronal excitability in response to glutamate stimulation in the secondary screens. This effect is observed intra-well, where sgRNA-containing neurons have a lower CaMPARI2 ratio than those that did not receive a sgRNA. Non-targeting controls (NTCs) are the same as in Figure 3A. Paired ratio t-test made between neuron populations receiving sgRNA (sgRNA+) and not receiving guide (sgRNA-). N = 3-6 independent culture wells per measurement. Each individual experiment was normalized to the excitability (CaMPARI2 response) of wells where neurons did not receive a sgRNA. For intra-well comparisons, a paired ratio t-test between sgRNA– and sgRNA+ populations were performed for each sgRNA. The calculated P values were corrected for multiple hypotheses (Holm-Šídák method). (b) CaMPARI2 neurons expressing sgRNAs targeting genes that, when knocked down, increased neuronal excitability in response to glutamate stimulation in the secondary screens. This effect is observed intra-well, where sgRNA containing neurons have a higher CaMPARI2 ratio than those that do not receive a sgRNA. Non-targeting controls (NTCs) are the same as in Figure 3A and Figure 5A. Paired ratio t-test made between neuron populations receiving sgRNA (sgRNA+) and not receiving guide (sgRNA-). N = 3-6 independent culture wells per measurement. Each individual experiment was normalized to the excitability (CaMPARI2 response) of wells where neurons did not receive a sgRNA. For intra-well comparisons, a paired ratio t-test between sgRNA– and sgRNA+ populations were performed for each sgRNA. The calculated P values were corrected for multiple hypotheses (Holm-Šídák method). (c) PICALM KOs exhibit a reduced rectification index (the ratio of current magnitude at +40 mV vs –70 mV) relative to NTC, suggesting an increased presence of synaptic calcium permeable AMPA receptors. The traces represent the mean and standard error of all sEPSC recordings from NTC and PICALM KO iNeurons and are shown normalized to the average –70 mV sEPSC for each neuron. (N = 15 NTC, 13 PICALM KO). (d) PICALM KOs displayed a decreased rectification index as compared to NTCs (N = 15 and 13 neurons for NTCs and PICALM KOs, respectively; analyzed using the Mann-Whitney test). Error bars donate standard error. (e) Sample sEPSC recordings from representative NTC and PICALM KO iNeurons.

We then examined three hits that increased neuronal excitability in the CaMPARI2 CRISPRi screens. *CACNG7* encodes TARP g-7, which is related to stargazing, has been found to be reduced in mouse models of spinocerebellar ataxia type 23 (SCA24).^57^ In the screen, *CACNG7* knockdown increased neuronal excitability and we observe a general increase in neuronal excitability in *CACNG7* knockdown neurons in independent experiments (Figure 5b). *PTEN*, classically associated with its role as a tumor suppressor gene^58^ and in mitophagy,^59^ has been shown to modulate neurite outgrowth and synapse density in *PTEN* KO mice.^60^ Furthermore, recent whole-exome sequencing studies have implicated *PTEN* in ASD.^15^ In the CaMPARI2 screen, *PTEN* knockdown increased synaptic excitability and *PTEN* knockdown neurons are more excitable when compared to baseline neurons (Figure 5b).

Recently, it has been suggested that *PICALM* may also regulate calcium permeable (CP)-AMPA receptors (AMPARs with homomeric GluA1 subunits) by targeting surface receptors for endocytosis.^61^ To test for regulation of CP-AMPARs in human iPSC-neurons, we performed patch clamp recordings in *PICALM* KO and non-targeting control iPSC-neurons. In these experiments we measured AMPA receptor mediated sESPC amplitudes at depolarized and hyperpolarized holding potentials and calculated the rectification index (sESPC_AMPAR_ +40 mV / sESPC_AMPAR_ –70 mV). While a high rectification index indicates few CP-AMPARs, a low rectification index indicates the presence of CP-AMPARs. We found a decrease in the rectification index in *PICALM* KO iPSC-neurons compared to non-targeting control (Figure 5c-e), suggesting that *PICALM KO* increases the level of CP homomeric GluA1 AMPARs at the synapses of human iPSC-derived neurons.

### CROP-seq screen connects transcriptional changes to excitability phenotypes

Many of the hits in the glutamate CaMPARI2 screen are known regulators of differentiation, neural development, and have been implicated in neurodevelopmental disorders. To evaluate which of these genes are hits because of changes to neuronal maturation in iPSC-neurons versus bona fide modulator of neuronal excitability, we generated a pooled sgRNA library targeting 45 genes (2 sgRNAs per gene, plus 5 non-targeting controls; SI table 10) that were annotated as being transcriptional regulators, involved in development or differentiation, and were hits in the CaMPARI screen. We introduced the CROP-seq library via lentiviral transduction into iPSCs at a low MOI (0.1) and selected transduced iPSCs with puromycin. These iPSCs were differentiated into neurons, dissociated into a single-cell suspension after 21 days of differentiation and subjected to single-cell RNA sequencing (scRNA-seq). After removing droplets that did not pass quality control cutoffs and only keeping cells with an sgRNA MOI of 1 (Methods), we analyzed 9252 neurons. By combining a transcriptional profile with assignment of a sgRNA for every neuron through sequencing, we could assess the effect of gene knockdowns in a pooled screen.

### Marker gene expression and cluster shift analysis does not indicate broad changes in cell identity

To examine the effect of knockdown on neuronal identity, we performed Leiden clustering on all 9252 neurons. By uniform manifold approximation and projection (UMAP) for dimensionality reduction, we separated cells into three clusters (Extended Data Figure 8a-e) and examined the expression of marker genes (Figure 6a). Doublecortin (*DCX*), a microtubule binding protein associated with the developing cortex, was highly expressed in all neurons. Eomesodermin (*EOMES*), a TBR1 family transcriptional activator critical for progenitor cell proliferation, and DNA topoisomerase II alpha (*TOP2A*), which is a marker for mitotically active cells, showed low expression throughout each cluster. For each neuronal and neural progenitor marker gene, expression was consistent across the defined clusters (Figure 6a), indicating that the clusters do not reflect different stages in neuronal differentiation.

**Figure 6.**
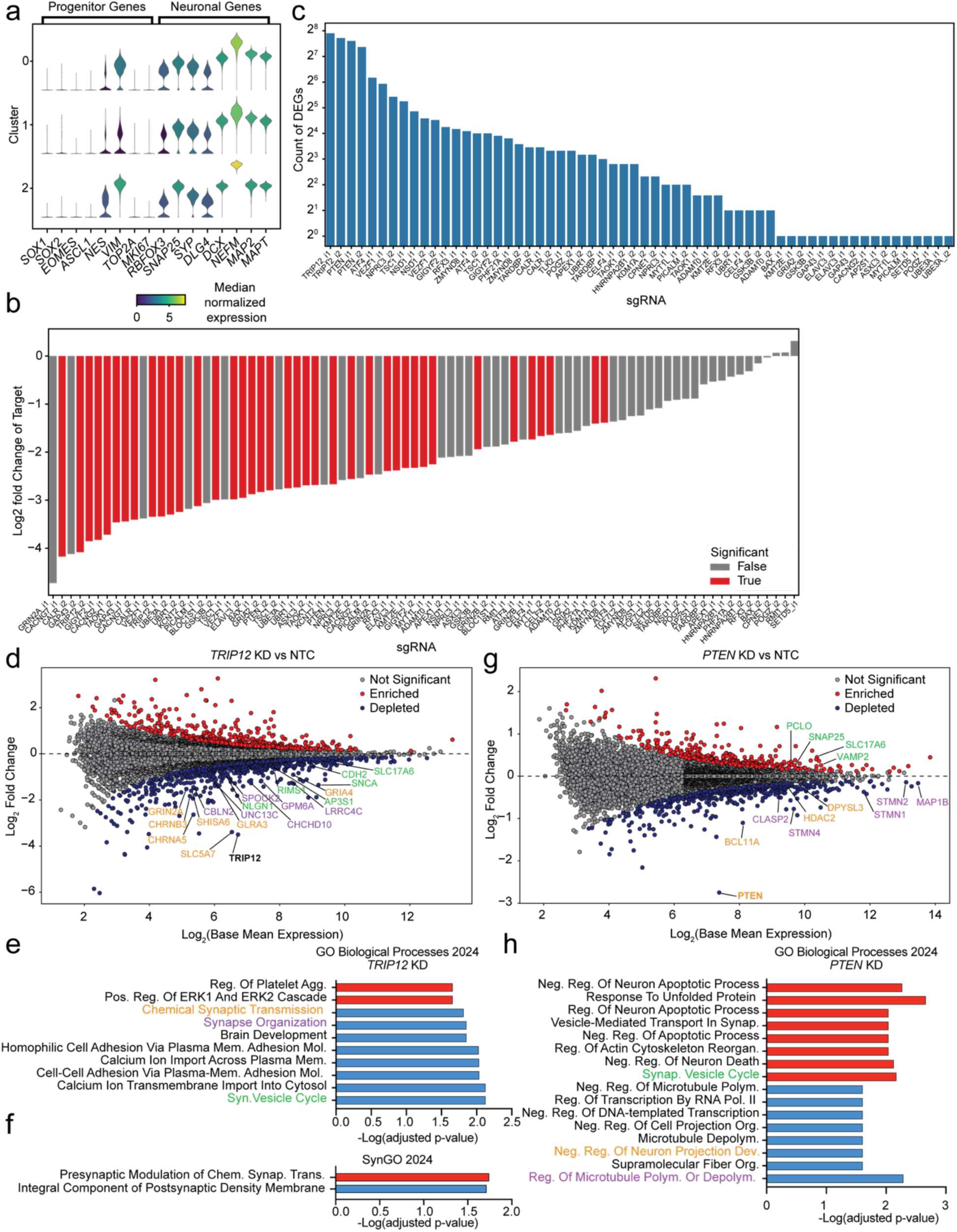
CROP-seq reveals transcriptional changes to genes involved in synaptic processes drive changes in excitability. (a) Marker gene expression is homogenous across clusters and suggests that neuronal identity is conserved. (b) On-target knockdown of target genes and the corresponding sgRNAs. Red bars indicated a significant KD in comparison to NTCs, while grey bars did not reach significance. (c) Number of significantly differentially expressed genes for each detected sgRNA. Each sgRNA was compared separately to the group of five non-targeting control (NTC) sgRNAs, and significance was determined at a threshold of and adjusted p-value <0.05. (d) MA plot of TRIP12 knockdown, showing genes (colors indicate associated GO term in Figure 6e) related to synaptic transmission and synapse organization. (e) GO term (Biological Processes) enrichment analysis of upregulated (red) and downregulated genes in TRIP12 KD (f) GO term (SynGO) enrichment analysis of upregulated (red) and downregulated genes in TRIP12 (g) MA plot of PTEN knockdown, showing an upregulation of synaptic processes and downregulation of microtubule organization and polymerization. Genes are colored ti indicate associated GO term in Figure 6h. (h) GO term (Biological Processes) enrichment analysis of upregulated (red) and downregulated genes in PTEN KD.

Next, we evaluated the effect of each knockdown on the distribution of neurons within the defined clusters relative to the distribution of neurons with non-targeting control sgRNAs. Of the 45 genes examined, only *TRIP12* knockdown showed consistent results across both sgRNAs tested. We observed a significant enrichment of neurons with *TRIP12* knockdown in cluster 2 and a depletion from cluster 0 (Extended Data Figure 8f).

We also observed significant cluster enrichment (*RFX3*, *UBR1*) and depletion (*GIGYF2*, *VEZF1*) for individual sgRNAs and large, but statistically insignificant, percent changes in cluster occupancy for additional targets. Together with the consistent marker gene expression across the identified clusters, this suggests that changes in neuronal differentiation trajectory were not the main contributor to the observed changes in neuronal excitability for the hits in the CaMPARI2 screens, but that the hit genes affect neuronal activity through other mechanisms.

### Differential Expression Analysis uncovers differences in transcriptional signatures

To determine the transcriptional changes that resulted from each knockdown, we next used the CROP-seq dataset to identify differentially expressed genes (DEGs) by comparing each individual sgRNA knockdown to the group of 5 NTC sgRNAs. 39 sgRNAs had statistically significant knockdown of the intended target gene relative to NTCs (Figure 6b), although almost all sgRNA showed a trend for knockdown of the target gene, suggesting that knockdown was effective for most sgRNA, even if its detection was limited by statistical power. 43 of the 90 sgRNAs had greater than one DEG (Figure 6c). We identified 660 unique DEGs, 196 of which were shared across knockdown of two or more targeted genes.

Next, we examined sgRNA groups with on-target knockdown of the target gene (Figure 6b) with at least 10 significantly expressed genes. We performed hierarchical clustering to determine any common DEG signatures across knockdowns and found that pairs of effective sgRNAs targeting the same gene clustered together (Extended Data Figure 8g). However, we did not observe any convergence between DEG signatures of different gene knockdowns, suggesting the knockdowns we selected are acting through different pathways.

### TRIP12 knockdown induces transcriptional changes to the synapse

*TRIP12* encodes thyroid hormone receptor interactor 12, an ubiquitin E3 ligase involved multiple ubiquitin-dependent proteolytic pathways^62^ and DNA damage repair.^63,64^ Mutations in *TRIP12* have been associated with Clark-Baraister syndrome and intellectual disability^64^, however it remains unclear which mechanisms lead to the clinical presentation of disease. Given that *TRIP12* knockdown reduced excitability in the CaMPARI2 screen, significantly changed cluster occupancy, and resulted in the most DEGs, we examined the affected pathways caused by *TRIP12* knockdown in our neurons.

*TRIP12* knockdown resulted in 213 downregulated and 124 upregulated genes (Figure 6d). We performed gene set enrichment analysis on the upregulated DEGs, which were significantly enriched for GO terms related to ERK1/2 signaling (Figure 6e) and presynaptic changes to synaptic transmission (Figure 6f). Downregulated DEGs showed a significant enrichment for many terms related to synaptic transmission and development (Figure 6e) as well as components of the postsynaptic density (Figure 6f). These changes to the post-synapse, in particular the downregulation of the ionotropic glutamate receptor subunits *GRIA4* and *GRIN2A*, could explain the reduced excitability we observed with *TRIP12* knockdown neurons when treated with glutamate in the CaMPARI2 screen.

### PTEN knockdown changes neuronal projection regulation and synaptic vesicle cycling

Finally, we examined *PTEN* knockdown, which increased synaptic excitability in the CaMPARI2 screen and in individual validation experiments (Figure 5b). We found a downregulation of the genes *STMN1*, *STMN2*, and *STMN4* (Figure 6g), which encode stathmins involved in the regulating microtubule polymerization and depolymerization. This could explain previous studies in *PTEN* KO mice, which have enhanced dendritic arborization and additional protrusions compared to control mice.^60^ Additionally, gene set enrichment analysis of downregulated genes significantly highlighted Gene Ontology terms related to neuronal projections and microtubule polymerization (Figure 6h). We also performed the same analysis on upregulated genes and found significant enrichment of components of the synaptic vesicle cycle. Specifically, the upregulation of *SNAP25, VAMP2, PCLO,* and *SLC17A6* suggests changes in synaptic vesicle trafficking and regulation of SNARE complexes. This could explain both the increased excitability of *PTEN* knockdown neurons in the CaMPARI screen and the increase in frequency of excitatory quanta in *PTEN* KO mice from previous studies.^60^

## Discussion

Changes in neuronal excitability and synaptic dysfunction appear in many neurodegenerative^34,41,65–67^ and neurodevelopmental^35,38,68^ disorders. In this work, we developed a screening paradigm using calcium integration as a proxy for neuronal activity to explore how perturbation of genes known to regulate to neuronal activity and/or identified risk genes affect synaptic excitability of iPSC-neurons. We found that we could detect the effect of glutamate receptor subunits, TARPs, and ion channels. These hits were confirmed with patch clamp studies and suggests that mechanisms such as altered depolarization-induced action potential dynamics, altered synaptic receptor kinetics, and changes in AMPA receptor calcium permeability could underlie the observed screen phenotypes. The screens and validation experiment also confirmed that roles in neuronal excitability for disease-linked genes could be detected with this approach.

Intriguingly, the CaMPARI2 signal in the primary FACS based screen proved to be a sensitive readout detecting a variety of changes in neuronal properties. This is even more remarkable given the heterogeneous response of individual iPSC-derived neurons to glutamate stimulation (Fig. 1f-h, Extended Data Fig. 2). These findings suggest that CaMPARI2-based CRISPR screens can detect modifiers of neuronal properties that are only present in a subset of the neurons screened, and validates CaMPARI2 as a suitable proxy to capture diverse neuronal phenotypes.

The CaMPARI2 screen evaluated the effect of known modifiers of synaptic excitability in iPSC-neurons. Multiple genes belonging to the transmembrane AMPA receptor protein (TARP) family were identified as hits. TARPs are known to influence the density and function of AMPARs.^49,54^ The type-I TARP encoded by *CACNG2* has previously been shown to interact with PSD-95 to stabilize AMPARs at the postsynaptic density^51^ and to modulate AMPA receptor levels ^50,54^ and kinetics.^54^ Of the six known TARP isoforms,^50^ CRISPRi knockdown of *CACNG2* and *CACNG4* reduced synaptic excitability while *CACNG7* knockdown increased synaptic activation in the CaMPARI2 based screen (Figure 2b).

Our results indicate that the CaMPARI2 screen is capable of detecting subtle changes to neuronal excitability and can highlight previously under-studied channels for additional characterization. *KCNT2* encodes the sodium-activated potassium channel K_Na_1.2 (Slick). Potassium (K^+^) channels play a major role in modifying neuronal excitability and regulating synaptic vesicle release,^69^ and K_Na_1.2 has been implicated in the adaptation of neuronal firing patterns in auditory neurons.^70^ Gain-of-function variants in the K_Na_1.1 (Slack) channel encoded by *KCNT1*, a paralogue of *KCNT2*, have been extensively linked to neurodevelopmental disorders (NDD) and epilepsy;^71,72^ however, the function of K_Na_1.2 is less understood, despite the connection of K_Na_1.2 variants to developmental and epileptic encephalopathy^73^ and epilepsy.^74^ In our study, knockdown of *KCNT2* increased excitability in iPSC-neurons in the CaMPARI2 screen (Figure 2b) and led to premature depolarization block in current-clamp experiments (Figure 3f-g). This depolarization block may lead to sustained activation of Ca^2+^ channels and lead to an increase in CaMPARI2 signal in our screen and flow cytometry experiments with 50% of neurons with *KCNT2* KD. At the same time, in cultures with 100% of neurons with *KCNT2* KO/KD, the reduced numbers of action potentials could lead to reduced network activity which may have a dominant effect in reducing overall excitability that overrides the cell-autonomous increases in calcium influx. This would lead to the decrease in CaMPARI signal we observed by flow cytometry in cultures where 100% of neurons have *KCNT2* KD. However, since we did not investigate the effects on synaptic excitability directly, we cannot rule out potential contributions *KCNT2* may have on synaptic excitability. This could be addressed in the future with additional electrophysiology studies.

Evaluating disease risk genes for their role in neuronal activity presents a possible route to developing new therapeutics. *PICALM* encodes the phosphatidylinositol-binding clathrin assembly protein (PICALM), and is originally noted for its involvement in AP2-dependent clathrin-mediated endocytosis.^75,76^ GWAS studies have implicated variants at the *PICALM* locus to be associated with late-onset AD,^77,78^ which has been verified with additional meta-analysis^79,80^ and functionally shown to modify the clearance of tau.^81^ We observed an increase in synaptic excitability with *PICALM* knockdown in the CaMPARI2 screen with glutamate, and PICALM knockdown increased excitability with the largest effect size of the genes tested in the individual validation experiments (Figure 5b). Other studies have noted late-onset AD risk genes, including *PICALM*, to be involved with synaptic dysfunction^34^ and a recent study demonstrated the role of *PICALM* in the regulation of long-term potentiation and memory.^61^

To our knowledge, this calcium-integration strategy represents the first pooled modifier screen for neuronal activity. Intriguingly, this strategy uncovered modifiers of numerous aspects of neuronal excitability, including depolarization block, changes to receptor kinetics, and modifications to receptor subunit partitioning at the synapse. Additionally, combining this method with scRNA-seq methods such as CROP-seq identified multiple genes with effects on excitability through changes to transcriptional programs of synaptic genes and neurite growth. Paired together, these complementary methods can identify genetic regulators of neuronal excitability and prioritize them for additional study.

While these CRISPRi screens using CaMPARI2 can uncover potentially novel roles for genes in neuronal activity and excitability, we note there are limitations with this method. Primarily, the 405 nm (UV) light needed for CaMPARI2 photoconversion can cause photodamage to cells. We demonstrate that neurons exposed to the five minutes of 405 nm light used in the screen do not undergo massive death or enter apoptosis on the same timescale of the screen, however we acknowledge low cell recovery from FACS, which we attribute to the fragility of these neurons. We were able to overcome this obstacle because of the scalability of our iPSC-neuron system. However, screening the entire protein coding genome will be a major endeavor. To combat this drawback, we plan to incorporate other calcium integration strategies such as scFLARE.^82^ This strategy uses 488 nm light to initiate activation through a conformational change of the sensor, which is less toxic than UV light. Furthermore, this Ca^2+^ integration strategy can be modified to provide a transcriptional readout,^83^ which could be paired with CRISPRi for transcriptional perturbation screens using either the Perturb-seq or CROP-seq method.

Another potential drawback of our first application of this approach is the limited functional maturity of iPSC-neurons. We chose to perform these studies after 21 days of differentiation in BrainPhys media, which helps to accelerate maturation of iPSC-derived cultures,^84,85^ as this provided a timepoint where the neurons displayed reasonable spontaneous activity while also convenient for screening. Generally, extended maturation times upwards of 9 weeks (>60 days) are needed to develop synaptic maturity in these iPSC-neuron systems^86^ and co-culture with astrocytes is needed to enhance the maturation of iPSC-neurons and facilitate increases in activity.^87^

In the first application of this CaMPARI2-based pooled screen, we decided focus on synaptic excitability due to prevalence of synaptic defects in many neurological disorders. However, the low correlation between glutamate and KCl screens suggests that this method could be used to detect excitability modifiers through different mechanism. For this reason, it would be informative to perform additional large-scale screens to screen additional mechanisms of neuronal excitability. Furthermore, this screening approach could be adapted to examine neuronal activity in response to other pharmacologic agents like kainic acid to produce seizure-like firing. CaMPARI2 screens could also be used to evaluate electrophysiological maturation of iPSC-neurons longitudinally or to probe mechanisms of plasticity. Optogenetic tools that are spectrally compatible with CaMPARI2 could be used in this paradigm to examine how particular genes affect specific firing patterns of neurons.

We believe that our CaMPARI2-based CRISPRi screening platform will help to uncover the functional roles of risk genes for many different neurological disorders of both the central and peripheral nervous systems. The development of new differentiation protocols and increased availability of iPSC models provides an opportunity to extend this paradigm to other specialized types of neurons, such as motor neurons and inhibitory neurons, and to patient-derived iPSC-neurons with disease-linked mutations. Beyond astrocytes, other cell types could also be incorporated into 2D or 3D culture systems, such as iAssembloids.^27^ Understanding the genetic determinants of neuronal activity in different contexts will lead to an understanding of disease mechanisms related to aberrant neuronal activity potential and the development of new therapeutics.

## Methods

### Human iPSC Culture

hiPSCs were cultured in StemFlex medium (Thermo Fisher Scientific, A3349401) in plates or dishes coated with Growth Factor Reduced, Phenol Red-Free, LDEV-Free Matrigel Basement Membrane Matrix (Corning, 356231) diluted 1:100 in Knockout DMEM (Thermo Fisher Scientific, 10829–018). StemFlex medium was replaced daily. Upon 70-80% confluence, hiPSCs were dissociated to single-cell suspensions with StemPro Accutase Cell Dissociation Reagent (Thermo Fisher Scientific, A1110501). hiPSCs were washed with Dulbecco′s Phosphate Buffered Saline without CaCl_2_ and MgCl_2_ (Sigma, D8537-24X500ML), incubated with Accutase for 3-7 minutes at 37°C until cell could be dislodged with gentle trituration. The cell suspension was diluted 5x with DPBS, transferred to tubes, and centrifuged at 200xg for 5 minutes. hiPSCs were resuspended in StemFlex Medium supplemented with 10 nM Y-27632 dihydrochloride ROCK inhibitor (Tocris Bioscience, 1254), counted, and plated onto Matrigel-coated plates or dishes. hiPSCs were maintained in Y-27632 until colonies reached the appropriate size (more than 15 cells/colony). Studies with human iPSCs at UCSF were approved by the Human Gamete, Embryo, and Stem Cell Research (GESCR) Committee. Informed consent was obtained from the human subjects when the hiPSC lines were originally derived.

### Generation of CaMPARI2 hiPSC line

To generate CaMPARI2 iNeurons, the CaMPARI2 gene (1446 bp) was subcloned from pAAV_hsyn_NES-his-CaMPARI2-WPRE-SV40 (Addgene 101060, gift from Eric Schrieter)^32^ into a lentiviral plasmid (pRT050) under a CAG promoter and an upstream UCOE element via Gibson Assembly to yield pRT074. iPSCs expressing CRISPRi machinery and inducible NGN2^23^ were transduced with lentivirus delivering pRT074 (described in more detail below). Clonal lines were isolated by colony picking, differentiated to neurons, and evaluated for homogeneity of CaMPARI2 expression and dynamic range of photoconversion. A clonal line satisfying the above criteria was selected for CaMPARI2 photoconversion experiments and screening.

### Lentiviral transduction of iPSCs

#### CaMPARI2 (pRT074)

pRT074 was lentivirally packaged in HEK 293T cells (ATCC Cat. No. CRL-3216) DMEM complete medium: DMEM (Gibco, 11965-092) supplemented with 10% FBS (VWR, 89510, lot: 184B19), 1% penicillin-streptomycin (Gibco, 15140122), and 1% GlutaMAX (Gibco, 35050061) as described previously,^23^ using TransIT Lenti Transfection Reagent (Mirus Bio; Cat. No. MIR 6600) according to manufacturer’s protocol.

#### sgRNA constructs

Individual or pooled sgRNAs were lentivirally packaged in HEK 293T cells (ATCC, CRL-3216) in DMEM complete medium as described previously^23^ with the following modifications: a 3:1 ratio of 1 µg/µl polyethenylamine (PEI) (Linear, MW 25,000, Polysciences, 23966) to transfer DNA was used in place of TransIT Lenti Transfection Reagent. Cells were selected with 0.8 μg/mL puromycin (Gibco; Cat. No. A11138-03) for 2 days until the fraction of infected cells was > 0.95, as determined by flow cytometry of sgRNA-BFP fluorescence. Additional rounds of puromycin selection were used as necessary until the fraction of infected cells was above 0.95, after which cells were cultured for a single passage in the absence of puromycin to allow them to recover. sgRNA protospacer sequences are provided in Supplementary Table 7.

### iNeuron differentiation and culture

#### For screening, flow cytometry, and imaging

hiPSCs were differentiated into iNeurons according to Tian et al.^23^ At day 0 of differentiation, iNeurons were plated onto poly-D-lysine-coated plates (Corning 6W 356413, 12W 356470, 24W 354414, 96W 354640, 15cm 354550) in BrainPhys neuronal media (composed according to Bardy et al. 2015)^84^ and 2 µg/ml doxycycline (Takara 631311). Three days after plating and, later, twice weekly, half of the culture media was removed and replenished with freshly supplemented BrainPhys without doxycycline. iNeurons were cultured for 21 days for all activity experiments.

#### For electrophysiology with wild-type NGN2 iNeurons or RNP-CRISPR mediated knockouts (KO)

Human iPSCs from iP11N background (ALSTEM) with inducible expression of Ngn2 by doxycycline (Dox) were differentiated into neurons using combined Ngn2 programming and dual SMAD patterning according to optimized from a previously published protocol.^88,89^ The iPSCs were cultured in mTeSR plus media for maintenance and in initiation media (F12/DMEM media, N2, B27 without VA, glutamax, NEAA) for differentiation. On day 7 of differentiation, Ngn2 iNeurons were further cultured in neuron maturation medium (iCell Neural Base medium 1 CDI M1010, iCell Neural Supplement B CDI M1029, iCell Nervous system Supplement CDI M1031) for more than 14 days before electrophysiology measurements were made.

### iAstrocyte differentiation and culture

Neural progenitor cells (NPCs) were first differentiated as previously described,^26^ then iAstrocyte differentiation was initiated by seeding NPCs into ScienCell Astrocyte Media (ScienCell Research Laboratories, 1801) + 2 µg mL^−1^ doxycyline.^26^ Media was changed every other day, with doxycycline maintained at 2 µg mL^−1^ throughout the differentiation process. At day 15, iAstrocytes were cryopreserved in ScienCell Astrocyte Media + 10% DMSO.

### Coculture of iNeurons and iAstrocytes

At day 0 of the iNeurons differentiation, day 15 iAstrocytes were thawed and resuspended in ScienCell Astrocyte media. iNeurons were dissociated and resuspended in BrainPhys neuronal media (see above). At a final seeding density of 2:1 iNeurons:iAstrocytes (468k iNeurons + 234k iAstrocyte per cm^2^), these cells were mixed in a 1:1 ratio of BrainPhys neuronal media: ScienCell Astrocyte media + 2 µg mL^−1^ doxycycline and seeded onto PDL + laminin coated dishes. After 3 days, a full media change into BrainPhys neuronal media was performed, followed by a half change with BrainPhys neuronal media every other day until the experimental timepoint.

For flow cytometry and FACS, these cultures were dissociated with the same method as monocultured neurons (see above), and neurons were gated using the CaMPARI green fluorescence.

### RNP-CRISPR knockouts

On day 7 of differentiation, cells were lifted by Accutase (Stemcell technology) and resuspended in P3 buffer (Lonza). RNP were prepared by mixing Cas9 (Alt-R™ S.p. Cas9 Nuclease V3, IDT) and gRNA for *KCNT2*, *CACNG2*, *PICALM* (Synthego KO kit) and non-targeting control (NTC) sequences (Genentech) separately. KO kits were used as a pool of three sgRNAs/gene (protospacer sequences are provided in Supplementary Table 6). These were then incubated at room temperature for 20 min. iNeurons in P3 buffer were then mixed with Cas9 RNP and nucleofected using pulse program CV-110 in the 4D-Nucleofector (Lonza). After nucleofection, iNeurons were plated at 1M in poly-D-lysine (A38904-01, Gibco) and iMatrix (T304, TaKaRa) coated µ-dishes (Cat. No. 80136, ibidi) for electrophysiology and further cultured in neuron maturation medium (iCell Neural Base medium 1 CDI M1010, iCell Neural Supplement B CDI M1029, iCell Nervous system Supplement CDI M1031) for more than 14 days before electrophysiology measurements were made.

### iNeuron dissociation protocol for flow cytometry

Dissociation solution was made from resuspending papain (Worthington; Code: PAP2; Cat. No.LK003178) in 1X Hank’s Balanced Salt Solution (Corning; Cat. No. 21-022-CV) to 20 U/mL. 5 mM MgCl_2_ and 10 U/mL DNaseI (Worthington; Code: DPRF; Cat. No. LS006333) were then added to the solution. Dissociating solution was added to 96-well, 24-well, 12-well, 6-well plates and 15 cm dishes at volumes of 20 µL, 100 µL, 250 µL, 500 µL, and 5 mL, respectively. iNeurons were treated with dissociation solution for 12 minutes at 37°C and resuspended in 1X Hank’s Balanced Salt Solution (HBSS, Gibco) before analysis.

### CaMPARI2 photoconversion

CaMPARI2 iNeurons plated on poly-D-lysine coated cell culture plates were illuminated with 405 nm light (LED output 39 W) in a temperature-controlled box with rotating turntable (FormLabs FH-CU-01). After 5 minutes of illumination at 37°C, iNeurons were dissociated (see above) or imaged.

For imaging CaMPARI2 photoconversion, CaMPARI2 iNeurons were plated onto 96-well poly-D-lysine coated plates (Corning 354640) at 25,000 cells/well (78,000 cells/cm^2^) in BrainPhys neuronal media. On day 21, iNeurons were treated with 30 µM glutamate, 1.5 mM tetrodotoxin, or 0.1% DMSO (Vehicle) for 5 minutes at 37 °C. After photoconversion (see above), iNeurons were imaged on an ImageXpress Micro Confocal High-Content Imaging System (Molecular Devices) using an Apo LWD 20X/0.95NA water immersion objective. Laser settings: Hoechst 33342 – EX 405/20 nm, EM 452/45 nm with FF409-Di03 dichroic; CaMPARI2-green – EX 467/21 nm; EM 520/40 nm with ZT 405/470/555/640/730 dichroic; CaMPARI2-red – EX 555 nm, EM 598/40 nm with ZT 405/470/555/640/730 dichroic.

### CaMPARI2 photoconversion experiments for flow cytometry

CaMPARI2 iNeurons were plated onto 96-well poly-D-lysine coated plates (Corning 354640) at 25,000 cells/well (78,000 cells/cm^2^) in BrainPhys neuronal media. On day 21, iNeurons were treated with 30 µM glutamate, 1.5 mM tetrodotoxin, or 0.1% DMSO for 5 minutes at 37 °C. iNeurons were then photoconverted with UV light and dissociated as described above before analyzing them by flow cytometry using a BD LSRFortessa X14 (BD Biosciences) using BD FACSDiva (v.8.0.1.1) software. Flow cytometry data was analyzed using FlowJo analysis software (v10.10.0; BD Life Sciences; Ashland, OR, USA), R/G ratio was calculated for each event captured as the ratio of FITC and mCherry. Mean R/G was calculated for subsequent analyses of CaMPARI2 photoconversion.

### Toxicity assays

CaMPARI2 cells at Day 0 of differentiation were plated in triplicate at 80,000 cells per well onto 24-well (42k cells/cm^2^) poly-D-lysine-coated cellware (Corning 354414) in 400 µl/well of supplemented BrainPhys media. After 21 days of differentiation, CaMPARI iNeurons were UV illuminated for 0, 1, 5, 15, and 60 minutes. Viability was analyzed immediately after and 24 hours after UV exposure. CaMPARI2 iNeurons were dissociated as above, pelleted at 300xg for 5 minutes, and resuspended in 20 µl 1X HBSS, combined with 20 µL 0.4% Trypan Blue (Invitrogen T10282), and counted on a Countess II automated cell counter (Invitrogen AMQAX1000) to quantify viable cells.

CaMPARI2 iNeurons were plated onto 96-well poly-D-lysine coated plates (Corning 354640) at 25,000 cells/well (78,000 cells/cm^2^) in BrainPhys neuronal media. On day 21, these were illuminated with UV for 0, 1, 5, 15, or 30 minutes. To perform viability staining with TO-PRO-3 (ThermoFisher T3605), CaMPARI2 iNeurons were washed 3x with DPBS, stained with TO-PRO-3 diluted 1:1000 in DPBS and 1 µM Hoechst 33342 (Thermo Fisher 62249) nuclear counterstain for 30 minutes at room temperature. After staining, cells were washed 3x with DPBS and imaged on an ImageXpress Micro Confocal High-Content Imaging System (Molecular Devices) using an Apo LWD 20X/0.95NA water immersion objective. Laser settings: TO-PRO-3; EX 638/17 nm, EM 624/40 nm with ZT 405/470/555/640/730 dichroic; Hoechst 33342; EX 405/20 nm, EM 452/45 nm with FF409-Di03 dichroic. Images were analyzed and cells were segmented using CellProfiler (v 4.2.4) analysis software. TO-PRO-3 staining intensity per object was measured and difference in signal between TO-PRO-3 stained versus unstained cells provided basis for live/dead cell filtering. After filtering live and dead cells, the proportion of live cells was used for viability analysis.

### Glutamate titration

After 21 days of differentiation, CaMPARI2 iNeurons were treated with increasing concentrations of glutamate (0-500 µM) for 5 minutes. This was followed by CaMPARI photoconversion and dissociation (protocols as described above). The mean R/G ratio for each concentration, with 6 independent culture wells per measurement, was used to calculate a polynomial fit using Prism (v10.1.1). 30 µM was chosen for subsequent experiments and screens due to the potential dynamic range available to detect increases in activity (increase CaMPARI2 response) while still allowing for some dynamic range to sense decreases in activity (lower CaMPARI2 response).

### CRISPR screens

#### Generation of sgRNA library

A custom CRISPRi library targeting genes associated neurological disease and activity (Neuron Activity library) was synthesized as follows: The disease-associated gene list were generated using DisGeNET^90^ online database, using the neurodegenerative, neurodevelopment, and neuropsychiatric diseases highlighted in Figure 2c as search terms. This was supplemented using genes chosen by searching terms ‘synapse’, ‘neurotransmitter’, ‘calcium’, ‘membrane potential’, ‘neuron activity’, and ‘neuron membrane’ in GO, KEGG, Reactome and Wikipathways. This gene list was filtered for gene expressed in NGN2-iNeurons based of RNA sequencing data,^23^ resulting in a final number of 1343 genes targeted in the library. The top 5 predicted sgRNAs^91^ oligonucleotide sequences per gene plus 250 non-targeting control (NTC) sequences (for 7450 total library elements) were synthesized by Agilent Technologies (RRID:SCR_013575; Santa Clara, CA) on a 7.5K oligo array and cloned into the pLG15 sgRNA expression vector. Library complexity and the presence of each oligo was confirmed with Illumina next-generation sequencing.

#### Primary FACS based screen

The Neuron Activity library was packaged into lentivirus as previously described for the CRISPRi genome-wide sublibraries.^23^ CRISPRi-NGN2-CaMPARI2 iPSCs were infected by the sgRNA library at MOIs between 0.3-0.5 (as measured by the BFP fluorescence in the sgRNA lentiviral vector) with >1000X coverage per library element. On the next passage, iPSCs expressing the sgRNA vector were selected using puromycin (see above) as they were expanded for screening, limiting the total number of passages from lentiviral transduction to three to limit dropout of sgRNAs at the iPSC stage. The sgRNA library has a puromycin resistance gene driven from the strong EF1alpha promoter and confers resistance to high levels of puromycin compared to the low levels of expression of this gene from the NGN2 cassette that transcribed as part of the endogenous AAVS1 locus. This enables us to re-use puromycin resistance for selection of iPSCs transduced with the sgRNA construct. iPSCs were scaled to a cell count corresponding to >10e^4^ x coverage per library element for each FACS sorted population, and differentiated into neurons as described above. On day 0 of the differentiation, iNeurons were plated onto 15 cm poly-D-lysine-coated plates (Corning 354550) at a density of 138k/cm^2^. iNeurons were maintained as described above until FACS sorting.

On day 21, iNeurons were treated with 30 µM glutamate for 5 minutes before photoconversion (described above). These were then dissociated with the above dissociation protocol, with the following modifications: incubation time in dissociation solution was increased to 20 minutes, papain was resuspended in 1:1 HBSS and Accutase, and dissociated neurons were resuspended in FACS wash buffer: 20 mL HBSS supplemented with 5 mM MgCl_2_, 10 nM Y-27632, and 10 U/mL DNaseI. Resuspended iNeurons were maintained as a sheet, transferred to a 50 mL tube, vigorously triturated, then pelleted at 300xg for 10 minutes. After the supernatant was discarded, the pellet was resuspended in 1-2 mL of FACS wash buffer, using trituration to make a single-cell suspension. Resuspended iNeurons were sorted into high and low R/G ratio populations corresponding to the top 35% and bottom 35% of the R/G signal distribution (see gating strategy, Supplementary Information), followed by sample preparation for next-generation sequencing. This screen was conducted in duplicate.

#### Pooled secondary screens

The focused secondary screening library contained 1211 sgRNAs targeting 411 genes (3-5 sgRNAs per gene) that were hits in either of the replicate primary screens and 250 NTC sgRNAs (1622 total sgRNAs). sgRNAs were selected based on their phenotype in the primary screens. A pool of sgRNA-containing oligonucleotides were synthesized by Twist Biosciences (South San Francisco, CA, USA) and cloned into an optimized sgRNA lentivirus vector (pMK1334) as previously described.^24^ CRISPRi-NGN2-CaMPARI2 iPSCs were transduced with lentivirus containing the secondary screening library, selected, and expanded as described for the primary screen above. A fraction of day 0 iNeurons were harvested and subjected to sample preparation for next-generation sequencing.

On day 21, iNeurons were treated with either 50 mM KCl or 30 µM glutamate, undergoing the same screening protocol as above.

Separately, CRISPRi-NGN2-CaMPARI2 iPSCs transduced with lentivirus were treated with 30 µM glutamate to match the same photoconversion conditions as day 21 iNeurons. These were then dissociated with Accutase (see Human iPSC Culture subsection) and sorted into high and low R/G ratio populations corresponding to the top 35% and bottom 35% of the R/G signal distribution, followed by sample preparation for next-generation sequencing.

### Screen sample preparation

For each screen sample, genomic DNA was isolated using a Macherey-Nagel Nucleospin Blood kit (Macherey-Nagel; Cat. No. 740951.50) or Macherey-Nagel Nucleospin Blood L kit (Macherey-Nagel; Cat. No. 740954.20), depending on the number of cells recovered from FACS. sgRNA-encoding regions were amplified and sequenced on an Illumina HiSeq-4000 as previously described.^23^

### Validation of CRISPR screen hits

CaMPARI2 iNeurons expressing sgRNAs targeting hit genes were plated onto 96-well poly-D-lysine coated plates (Corning 354640) at 12,500 cells/well in BrainPhys neuronal media. The assay was internally controlled by plating an equal mixture of CaMPARI2 iNeurons expressing no sgRNA and knockdown iNeurons, to a final density of 25,000 cells/well (78,000 cells/cm^2^).

### qPCR

To quantify knockdown in CaMPARI2 cells, lysed cell pellets from human iPSCs or iNeurons were thawed on ice, and total RNA was extracted using Quick-RNA Miniprep Kit (Zymo; Cat. No. R1054). Complementary DNA was synthesized with the SensiFAST cDNA Synthesis Kit (Bioline; Cat. No. 65054). Samples were prepared for qPCR in technical triplicates in 10-µl reaction volumes using SensiFAST SYBR Lo-ROX 2X Master Mix (Bioline; Cat. No. BIO-94005), custom qPCR primers from Integrated DNA Technologies used at a final concentration of 0.2 µM and cDNA diluted at 1:3. qPCR was performed on an Applied Biosystems QuantStudio 6 Pro Real-Time PCR System using QuantStudio Real Time PCR software (v.1.3) with the following Fast 2-Step protocol: (1) 95 °C for 20 s; (2) 95 °C for 5 s (denaturation); (3) 60 °C for 20 s (annealing/extension); (4) repeat steps 2 and 3 for a total of 40 cycles; (5) 95 °C for 1 s; (6) ramp 1.92 °C s^−1^ from 60 °C to 95 °C to establish melting curve. Expression fold changes were calculated using the ΔΔCt method, and normalized to housekeeping gene *GAPDH*. Primer sequences are provided in Supplementary Table 5.

### Electrophysiology

Patch clamp electrophysiology measurements were made using a Multiclamp 700B amplifier, with pClamp10 acquisition software (Molecular Devices). Recordings were filtered at 2 kHz for voltage clamp or at 10 kHz for current clamp. Digitization was performed at a frequency of 10 kHz, with a Digidata 1440 analog-to-digital converter (Molecular Devices). For current clamp recordings the intracellular solution was composed of (in mM): 130 K-gluconate, 5 NaCl, 10 HEPES, 0.2 EGTA, 2 MgCl_2_, 4 Na-ATP, 0.5 Mg-GTP, with a pH of 7.3. For voltage clamp recordings, the intracellular solution was composed of (in mM): 130 CsMeSO_3_, 4 NaCl, 10 HEPES, 0.2 EGTA, 5 QX-314 Br, 4 Na-ATP, 0.5 Mg-GTP, with a pH of 7.3.

The recording chamber was continuously infused with oxygenated ACSF, which contained (in mM): 127 NaCl, 2.5 KCl, 1.25 NaHPO_4_, 25 Na_2_CO_3_, 25 glucose, 2.5 CaCl2, 1.3 MgCl2. Recording electrodes had an open-tip resistance between 4-8 MW. These recordings were performed 2-3 weeks after iNeurons culture media were switched to maturation medium or after the RNP-CRISPR knockout procedure. Data for each gene was collated from two batches of iNeuron cultures.

For examining effects of glutamate stimulation on iNeurons using electrophysiology, the current-spike output relationship was measured first as follows: Cells were current-clamped at –70 mV. A sequence of 20 x 1 second long current steps were injected, starting from –20 pA and increased by 10 pA for each step. Then, cells were recorded at their resting membrane potential and 30 µM glutamate in ACSF was washing in for 5 mins with flow speed adjusted to ∼2.5 ml/min to have recording chamber solution (∼4 ml) exchanged < 2 mins. Finally, cells were then subjected to perfused current-spike output relationship measurement again after perfusing aCSF for > 8 mins to monitor washout of glutamate.

### Specific electrophysiology protocols for individual validation of hit genes

For the *KCNT2* knockout and NTC iNeurons, the current-spike output relationship was measured as follows: Cells were current-clamped at –70 mV. A sequence of 20 x 1 second long current steps were injected, starting from –20 pA and increased by 10 pA for each step.

For *CACNG2* knockout and NTC iNeurons, spontaneous excitatory postsynaptic currents (sEPSC) were recorded for 1 minute while the cells were voltage clamped at –70 mV. In cases where no events were observed during the initial minute, an additional minute of recording was performed. To enhance the detection of synaptic events, the synaptic vesicle release probability was increased in these experiments by elevating KCl to 5 mM and CaCl2 to 4 mM with zero MgCl2 in ACSF.

For *PICALM* knockout and NTC iNeurons, cells were voltage clamped at –70 mV and +40 mV for one minute to measure sEPSCs. If no events were observed within the initial minute, the recordings were performed for an additional minute. AMPA receptor-mediated sEPSCs were isolated by adding either 50 µM D-AP5 or 100 µM DL-AP5 (Cat. No. 0106Tocris) to the recording solution. Addition of 100 µM cyclothiazide (Cat. No.0713, Tocris or Cat. No. ALX-550-338-M050 Enzo) was used to reduce AMPA receptor desensitization and facilitate the detection of synaptic events. 100 µM spermine (Cat. No. S3256 Sigma-Aldrich) was included in the intracellular solution to block calcium-permeable AMPA receptors at +40 mV, aiding in the quantification of the rectification index (mean sEPSC current amplitude at +40 mV / mean sEPSC current amplitude at – 70 mV).

All recordings were analyzed using either Easy Electrophysiology software (v2.6 or v2.7, Cambridge, UK) or Clampfit (v10.7, Molecular Devices; San Jose, CA, USA). The minimum current step that initiated action potential firing was defined as the rheobase. The maximum firing rate was calculated at the current step that initiated the highest number of action potentials, with no failure being observed. The action potential half-width was computed from the first action potential at the rheobase. Recordings were included for analysis if a minimum of 5 sEPSC events were detected during the recording time frame (0.08Hz). Charge transfer for each recording was computed as the frequency * 60 seconds * the charge transfer for the average sEPSCs. Statistical analyses were performed using GraphPad Prism (v10.1.1; GraphPad Software; Boston, MA, USA) including the Mann-Whitney test or an ANOVA, followed by the Kruskal-Wallis test for multiple comparisons.

### CROP-seq

#### CROP-seq sgRNA library construction

We selected genes that were hits in the CaMPARI2 screen that are known regulators of transcription and chromatin, plus control hit genes (e.g. *KCNT2*, *CACNG2*) for a total of 45 target genes. The top two performing sgRNAs from the CaMPARI2 screens were selected for each target. Five non-targeting control sgRNAs were added, resulting in a 95 sgRNA library (SI table 10).

Top and bottom oligo sequences for each sgRNA were ordered in an array from IDT, annealed in a 96 well plate, and then pooled. A 1:20 dilution of this pool was cloned into the CROP-seq vector pMK1334 via BstXI and BlpI restriction sites. To test ligation efficiency, a test transformation was performed in DH5alpha cells. This was followed by a large-scale transformation in Stellar Competent cells for final library amplification and purification. Before transducing cells, we measured the proportion of each sgRNA in the library through sgRNA enrichment PCR and next-generation sequencing to check sgRNA representation in the library and confirm a normal distribution.

#### Cell preparation

We produced a pool of lentiviruses containing the CROP-seq sgRNA library and transduced iPSCs at a low MOI (10% transduction efficiency) to minimize multiple infections in which a cell receives more than one sgRNA. Then, iPSCs were selected with puromycin (see above), differentiated into neurons, and grown to the same day 21 timepoint used for CaMPARI2 screen. These cells were dissociated in the same manner using papain, then pelleted at 300 xg for 5 minutes. These were resuspended and washed in dPBS. The centrifugation and wash steps were repeated for a total of 4 washes to completely remove papain, DNase, and cell debris. The resuspension was counted and diluted to target 20k cells for a single 10X Chromium GEM-X 3’ chip. Cells were prepared according to 10x Genomics general sample preparation protocol.

#### Single cell RNA-seq library preparation

20k cells were loaded into a single well of a 10X Chromium GEM-X 3’ chip for GEM preparation. Single cell library preparation was conducted according to the protocol provided for Chromium GEM-X Single Cell 3’ Reagent Kit v4 (10x Genomics, Cat#PN-1000686, User Guide CG000731 Rev A). Concentrations of libraries were measured using Agilent TapeStation D5000 reagents (Agilent Technologies, Cat#5067-5588) on an Agilent 4200 TapeStation system.

#### sgRNA enrichment

Briefly, three semi-nested PCR reactions were performed followed by 1x SPRIselect beads (Fisher Scientific, Cat#NC0406407) cleanup after each PCR reaction. In PCR1, 15ng full-length cDNA from single cell RNA-seq library prep per sample was used as a template. In PCR2, 10ng post-cleanup PCR1 product was used as a template. In PCR3, 10ng post-cleanup PCR2 product was used as a template to add sample indices. ll PCR reactions were conducted using KAPA Hotstart HiFi ReadMix (VWR, Cat#103568-584) with annealing temperature at 62°C (15 sec) and extension at 72°C (15 sec) for 18 cycles (PCR1) or 15 cycles (PCR2 and PCR3). Concentrations of post-PCR3 products were measured using Qubit dsDNA HS assay kit (Thermo Scientific, Cat#Q32854).

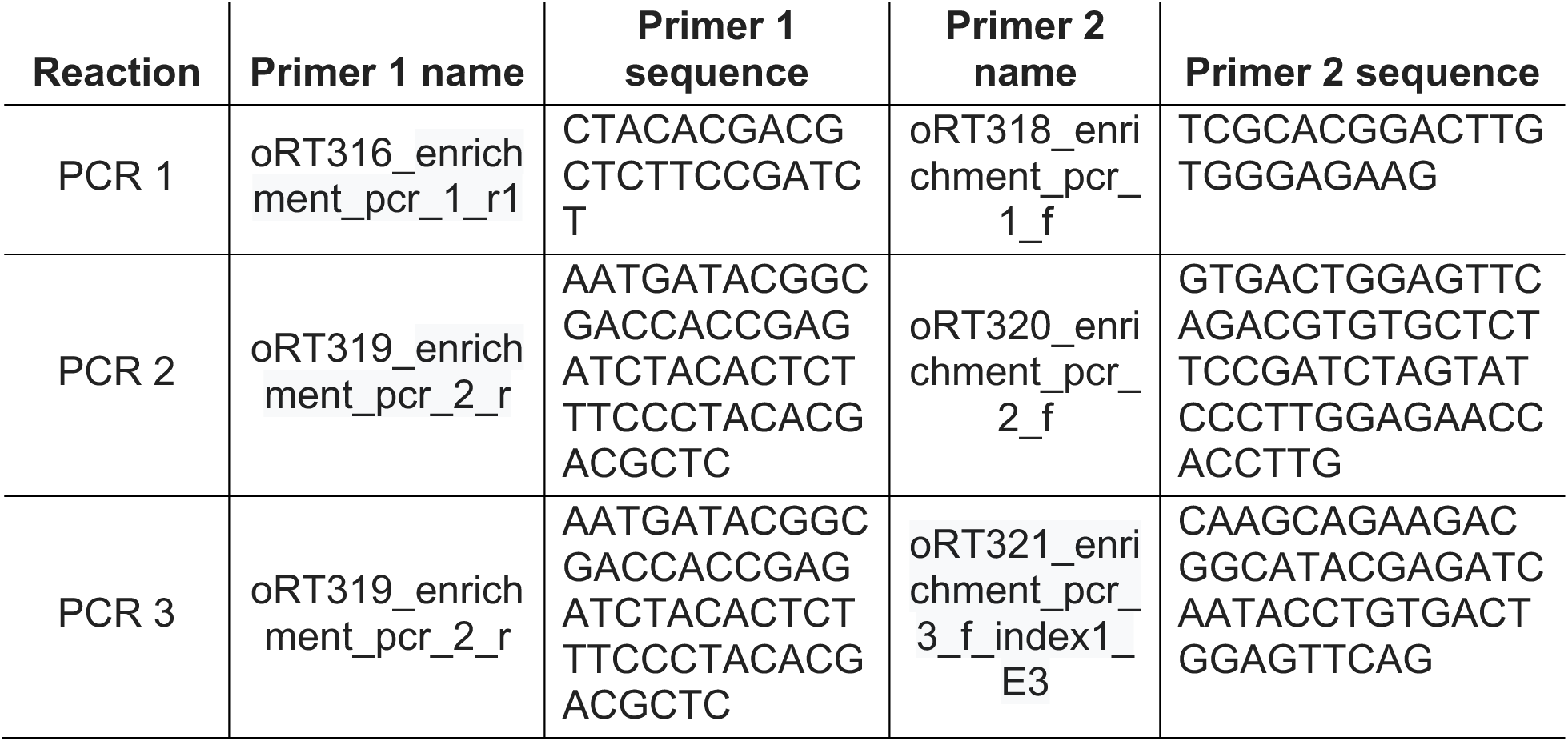

#### Sequencing

Samples of single cell RNA-seq libraries and enriched sgRNAs were pooled together at 5nM for sequencing on one lane of a 10B flow cell of an Illumina NovaSeqX following the recommendations from 10X Genomics. Approximately 1 billion and 400 million reads were generated for the gene expression and sgRNA enrichment libraries, respectively.

## Data Analysis

### CRISPR screen analysis

Primary screens were analyzed using sgcount and crispr_screen, two publicly available Rust packages based on MAGeCK^92^ and our previously published MAGeCK-iNC bioinformatics pipeline.^23^ The packages sgcount and crispr_screen are available on the Kampmann Lab website (https://kampmannlab.ucsf.edu/sgcount, https://kampmannlab.ucsf.edu/crisprscreen) and on Github (https://github.com/noamteyssier/sgcount, https://github.com/noamteyssier/crispr_screen).

Briefly, raw sequencing reads from next-generation sequencing are analyzed for the variable region (sgRNA protospacer sequence). Sequences in each sample are cropped to remove adapter regions and non-variable sequences to leave protospacer sequences remaining. These are then mapped to a sgRNA reference library to generate a count matrix for each screen sample. The quality of each screen was assessed by plotting the log_10_(counts) per sgRNA on a rank order plot using ggplot2.

Log_2_ fold change and significance P values were calculated for target genes over ‘negative-control-quasi-genes’, as well as an FDR given the ‘negative-control-quasi-genes’ that were generated by random sampling with replacement of five NTC sgRNAs from all NTC sgRNAs (iNC method). A ‘Gene Score’ was defined as the product of log_2_ fold change and −log10(P value). Hit genes were determined based on the p value cut-off corresponding to an empirical false discovery rate of 10%. Volcano plots of log_2_ fold change vs –log_10_(p value) and scatter plots of Gene Scores were generated using ggplot2.

### Over-representation analysis with WebGestalt

Hit genes from the primary screen were separated into hits decreasing and increasing activity. These lists were input into WebGestaltR^45^ (version 0.4.6) and enriched with the ORA method. The screening library was used as the reference gene list. Significant terms were thresholded at a FDR of 0.1, using Benjamini Hochberg.

### Image analysis with CellProfiler

Pipelines and example images are compiled in supplementary material, and all analyzes were performed using CellProfiler v.4.1.3. Cell toxicity by TO-PRO-3 co-localization: Nuclei were segmented as primary objects from Hoechst and TO-PRO-3 images. TO-PRO-3 objects that overlapped with Hoechst objects were counted as dead cells, while Hoechst objects negative for TO-PRO-3 were counted as live cells.

### Single cell RNA-sequencing alignment

#### Gene expression library sequence alignment

A kallisto index was generated from the human transcriptome using the ENSEMBL cDNA reference (GRcH38). Introns were annotated using the GRCh38 version 112 genome annotation. The default k-mer size (k=31) was used in generation of the index. The gene expression library sequences were then pseudo-aligned to this index using kallisto-bustools.^93^ After pseudo-alignment, barcodes were then corrected and filtered against the 10X-v4 cell barcode whitelist.

#### Knockdown Demultiplexing (sgRNA assignment)

A kallisto index was generated for the sgRNA library with a k-mer size of 15. The sgRNA enrichment PCR sequences were then pseudo-aligned to this index using kallisto bustools.^93^ After pseudo-alignment, barcodes were corrected and filtered against the 10X-v4 cell barcode whitelist. Cells were then assigned to guides using geomux (https://github.com/noamteyssier/geomux), which performs a hypergeometric test for each cell on its observed guide counts, then calculates a log2-odds ratio between the highest counts. Cells were assigned to their majority guide if their Benjamini-Hochberg corrected P-value was below 0.05, the log-odds ratio was above 1, and the total number of UMIs were greater than 5.

#### Single-Cell Analysis

##### Preprocessing

Cells were merged for the 10X sequence and the enrichment PCR by matching on cell-barcode and GEM library. Cells were filtered to only include those with a minimum count of 2000 and a minimum of 100 genes. 3000 of the most highly variable genes were selected with a minimum shared count of 10. Cells were further filtered to only include those with less and 8% and greater that 3% ribosomal gene content and less than 6.5% mitochondrial gene content. Cells were then normalized to a target sum of 1000 and log transformed. Principal component analysis was then performed using these highly variable genes. Clustering was performed using the Leiden algorithm, setting the resolution to 0.15. All single-cell filtering, transformation, and dimensionality reduction was performed using scanpy 1.10.3.

##### Differential Expression Analysis

Cells were grouped by knockdown, and GEM-library using ADPBulk (https://github.com/noamteyssier/adpbulk). Differential expression tests were performed using DESeq298 comparing each sgRNA knockdown against the non-targeting control sgRNAs.

##### Cluster overrepresentation analysis

We calculated Leiden cluster overrepresentation for each guide using a chi-square test for each guide/cluster in the dataset. Specifically, a chi-square test is performed for each knockdown group and Leiden cluster between each group distribution and each non-targeting control distribution. The p-values from this are then aggregated over the non-targeting controls using a geometric mean. Finally, these p-values are adjusted for multiple hypothesis testing using a Benjamini Hochberg correction. We performed this analysis using the open-source python module cshift (https://github.com/noamteyssier/cshift)

##### Knockdown Clustering Analysis and Gene Set Enrichment

Gene set enrichment analysis was performed using Enrichr and visualized using IDEA (https://github.com/noamteyssier/idea).

#### Software

GraphPad Prism v10.1.1; GraphPad Software; Boston, MA, USA

FlowJo analysis software v10.10.0; BD Life Sciences; Ashland, OR, USA

RStudio v2023.06.1+524 RStudio, PBC, Boston, MA, USA

Easy Electrophysiology software; v2.6; Cambridge, UK

Clampfit; v10.7; Molecular Devices; San Jose, CA, USA

*R packages used for presentation of data:*

tidyverse, ggrepel, ggpubr, reshape2, dplyr, ggplot2, grid, WebGestaltR, extrafont, viridis, RColorBrewer, ggthemes, remotes, circlize, corrplot, Hmisc.

#### Python (3.12) packages used for processing and presentation of scRNA-seq data

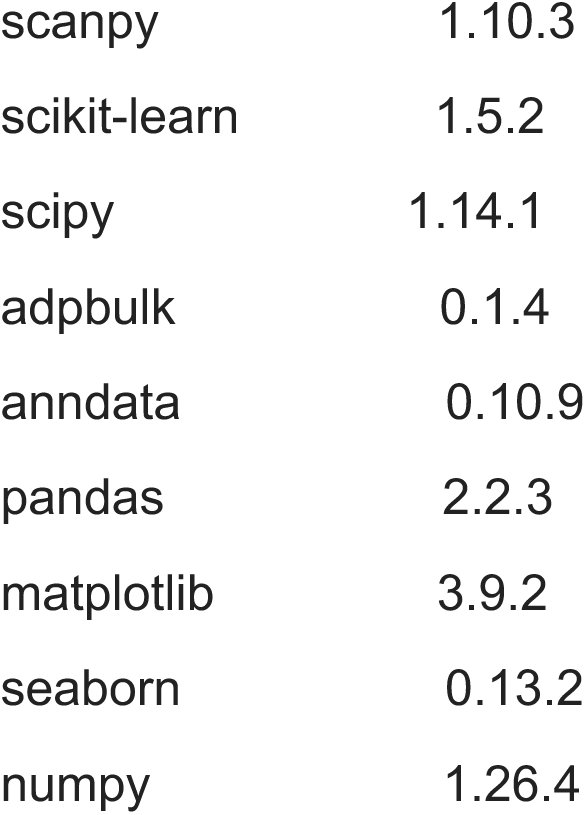

### Statistics and reproducibility

No statistical methods were used to pre-determine sample sizes but our sample sizes are similar to those reported in previous publications.^23–25^ No randomization of samples was used since treatment group of cells were generally derived from the same population of cells. Data collection and analysis were not performed blind to the conditions of the experiments. No data points were excluded from analysis. Data distribution was assumed to be normal but this was not formally tested.

### Reporting summary

Further information on research design is available in the Nature Research Reporting Summary linked to this article.

### Data availability

All screen datasets will be shared on the CRISPRbrain data commons (http://crisprbrain.org/) and will be shared upon request (associated with Figs. 2 and 4 and Extended Data Figs. 2). Single-cell RNA sequencing data will be deposited in NCBI GEO. There are no restrictions on data availability.

### Code availability

The sgcount and crispr_screen bioinformatics pipelines for analysis of pooled screens are available on the Kampmann Lab website (https://kampmannlab.ucsf.edu/sgcount, https://kampmannlab.ucsf.edu/crisprscreen) and on Github (https://github.com/noamteyssier/sgcount, https://github.com/noamteyssier/crispr_screen). Analysis pipelines used to analyze CROP-seq data are open source and available on Github (ADPBulk-https://github.com/noamteyssier/adpbulk; cshift-https://github.com/noamteyssier/cshift; IDEA-https://github.com/noamteyssier/idea).

The CellProfiler pipelines will be made available on request to the corresponding authors (M.K.) and will also be submitted to the CellProfiler depository of published pipelines (https://cellprofiler.org/examples/published_pipelines.html) upon publication.

## Availability of biological materials

All materials can be requested from the corresponding author (M.K.) and will be made available without restrictions via a material transfer agreement.

## Acknowledgements

We would like to thank Nick Page (UCSF) for contributions to preliminary experiments, the Weill Imaging Core and the Center for Advanced Light Microscopy, including Caroline Mrejen, So Yeon Kim, and Kari Herrington, for their technical support and microscopy instrumentation, the Laboratory for Cell Analysis and Sarah Elmes for use of FACS instruments.

We would like to thank Scott Martin, Ana Jovicic, and the Genentech Research and Early Development team (gRED) for their helpful discussions.

The authors would also like to thank Mor Alkaslasi, Olivia Teter and Avi Samelson for helpful comments and suggestions during the writing of this manuscript.

This research was funded by an award by the Alliance for Therapies in Neuroscience (ATN, Genentech and the University of California San Francisco) to M.K., an Alzheimer’s Association Research Fellowship to S.C.B. (23AARF-1027616), a California Institute for Regenerative Medicine (CIRM) grant to J.Y.C. (EDUC2-12730), a Rainwater Charitable Foundation Tau Consortium Investigator award, a Ben Barres Early Career Acceleration award by the Chan Zuckerberg Initiative, and NIH/NINDS grant U54 NS123746 to M.K.

Sequencing was performed at the UCSF CAT, supported by UCSF PBBR, RRP IMIA, and NIH 1S10OD028511-01 grants.

## Author contributions

Conceptualization of the CaMPARI2 based screen strategy was done by S.C.B, R.T, and M.K. CRISPRi screens and individual validation of screen hits were performed by S.C.B, V.G., E.M., and J.Y.C. sgRNA constructs for KD validation experiments were synthesized by S.C.B, V.G., J.Y.C, and L.Y. iNeuron-iAstrocyte coculture system was optimized by K.S. and S.C.B. CRISPR-RNP KOs in iPSC-neurons was performed by X. H., A.C, C.G.J., and C.E. Electrophysiology was performed by M.T. with input from J.H. CRISPR screen analysis was performed by S.C.B. with an analysis pipeline developed by N.T.. CROP-seq library preparation, cell preparation, and analysis was performed by S.C.B. and analysis was advised by N.T.. S.C.B, V.G., and M.T. created the figures, with input from all authors. S.C.B and M.K wrote the manuscript with input from all authors. All authors reviewed and approved the final manuscript.

## Figures

**Extended Data Figure 1.**
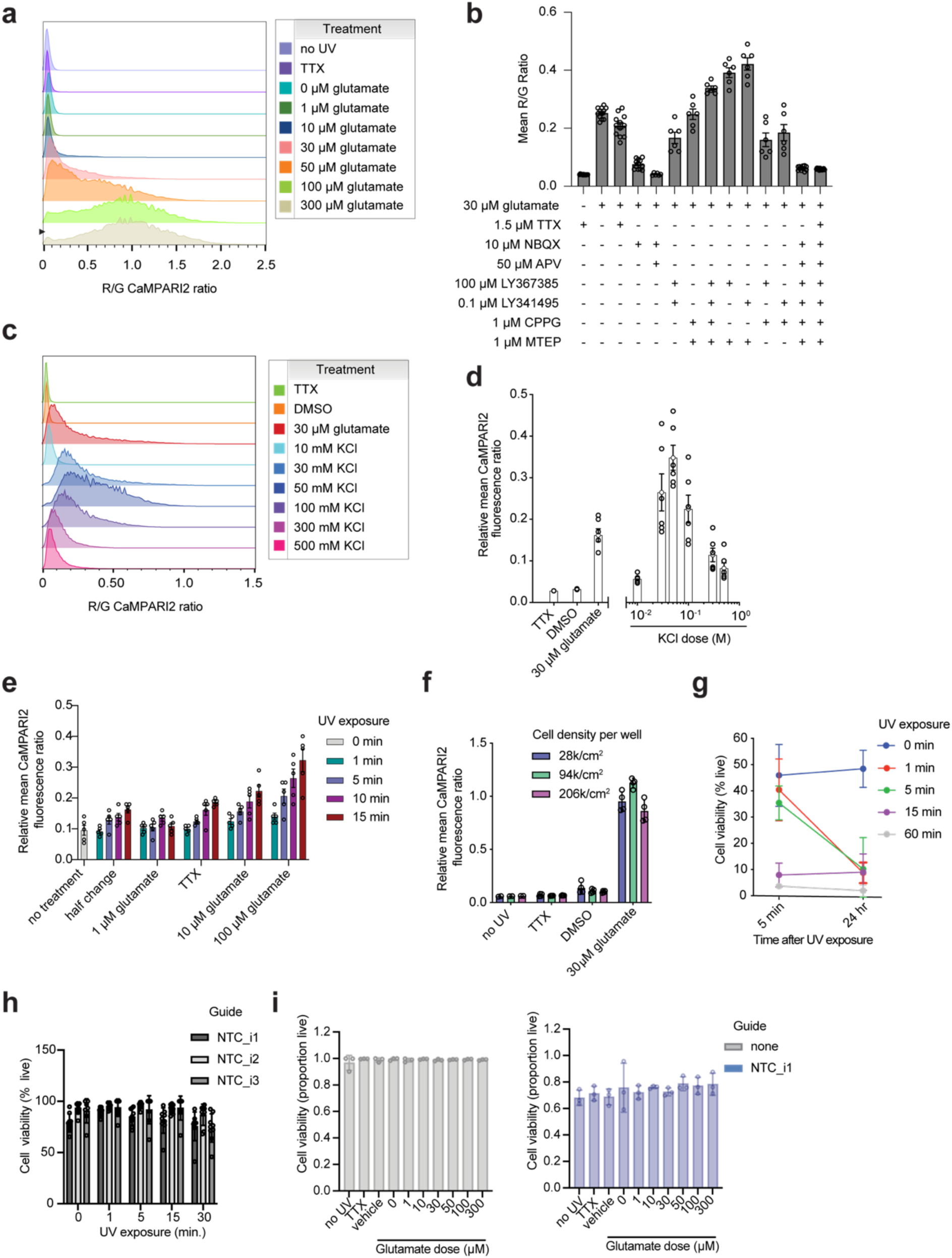
CaMPARI2 reliably conveys AMPAR activation at the synapse without significant UV toxicity. (a) CaMPARI2 dose response to glutamate. CaMPARI2 iNeurons were treated with TTX or various concentrations of glutamate (0, 1, 10, 30, 60, or 100 µM) and illuminated with UV to measure photoconversion of CaMPARI2. Photoconversion increased in a dose-dependent manner. (b) CaMPARI2 response to glutamate with co-treatment of GluR antagonists and TTX. CaMPARI2 activation is reduced to baseline (neurons treated with 1.5 µM TTX) when treated with APV and NBQX. Activation is not fully reduced to baseline with a full spectrum cocktail of mGluR antagonists, but is still lower than glutamate treatment alone. N = 6 independent culture wells. (c) CaMPARI2 dose response to KCl. CaMPARI2 iNeurons were treated with TTX, 30 µM glutamate, 0.1% DMSO or various concentrations of KCl (10, 30, 50, 100, 300, 500 mM) and illuminated with UV to measure photoconversion of CaMPARI2. Photoconversion increased in a dose-dependent manner to a peak at 50 mM, where a decline in photoconversion was observed at higher concentrations of KCl. (d) Quantification of CaMPARI2 red/green fluorescence ratio from (b). A non-linear fit could not be established, however the 50 mM KCl treatment was selected for later experiments due to the high conversion and range of response. N = 6 independent culture wells. (e) CaMPARI photoconversion in response to various UV durations. CaMPARI2 iNeurons were treated with TTX or various concentrations of glutamate (0, 1, 10, or 100 µM) and illuminated with UV for various durations of time (0, 1, 5, 10, or 15 min). After illumination, cells were dissociated for flow cytometry and analyzed. N = 5 independent culture wells. (f) Measuring effect of cell density on CaMPARI photoconversion. Significant deviations from optimal plating conditions had no significant effects on photoconversion upon TTX, vehicle, or glutamate treatment. N=4 independent culture wells. (g) Measuring UV toxicity in CaMPARI iNeurons. iNeurons were illuminated with UV for 0, 1, 5, 15, or 60 minutes and stained with Trypan Blue or TOPRO viability stain 5 minutes or 24 hours later to assess UV toxicity. N = 3 independent culture wells. (h) Measuring UV toxicity in CaMPARI iNeurons. CaMPARI iNeurons were illuminated with UV for 0, 1, 5, 15, or 30 minutes. Viability staining with TO-PRO-3 shows the proportion of viable cells after UV exposure. N = 9 independent culture wells. (i) Measuring glutamate toxicity. CaMPARI iNeurons were treated with 0, 1, 3, 10, 30, 50, 100 µM of glutamate for 5 minutes and UV illuminated for 5 minutes. TO-PRO-3 staining was performed to measure acute glutamate treatment toxicity. N = 3 independent culture wells.

**Extended Data Figure 2.**
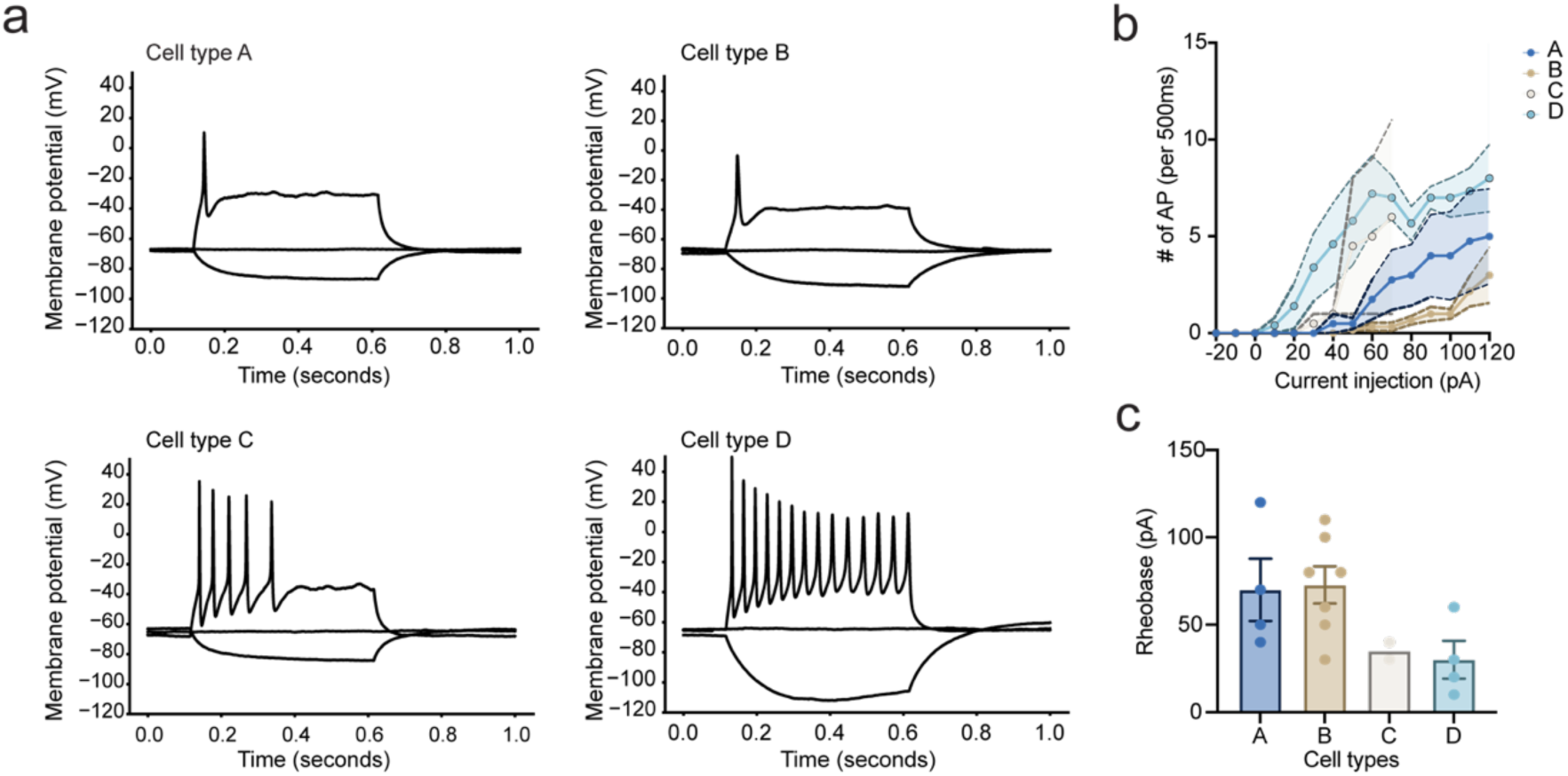
iNeurons have heterogeneous responses to glutamate stimulation. (a) Representative voltage responses from the current-spike output recordings of the corresponding iNeurons in Figure 1f-h. (b) Current-spike output relationships for all recorded iNeurons in Figure 1f-h. N = 2-7 independent neurons per cell type, 18 neurons total. Shaded error bars are S.E.M. (c) Rheobase values for all recorded iNeurons in Figure 1f-h. N = 2-7 independent neurons per cell type, 18 neurons total. One-way ANOVA, p = 0.05. Error bars are S.E.M.

**Extended Data Figure 3.**
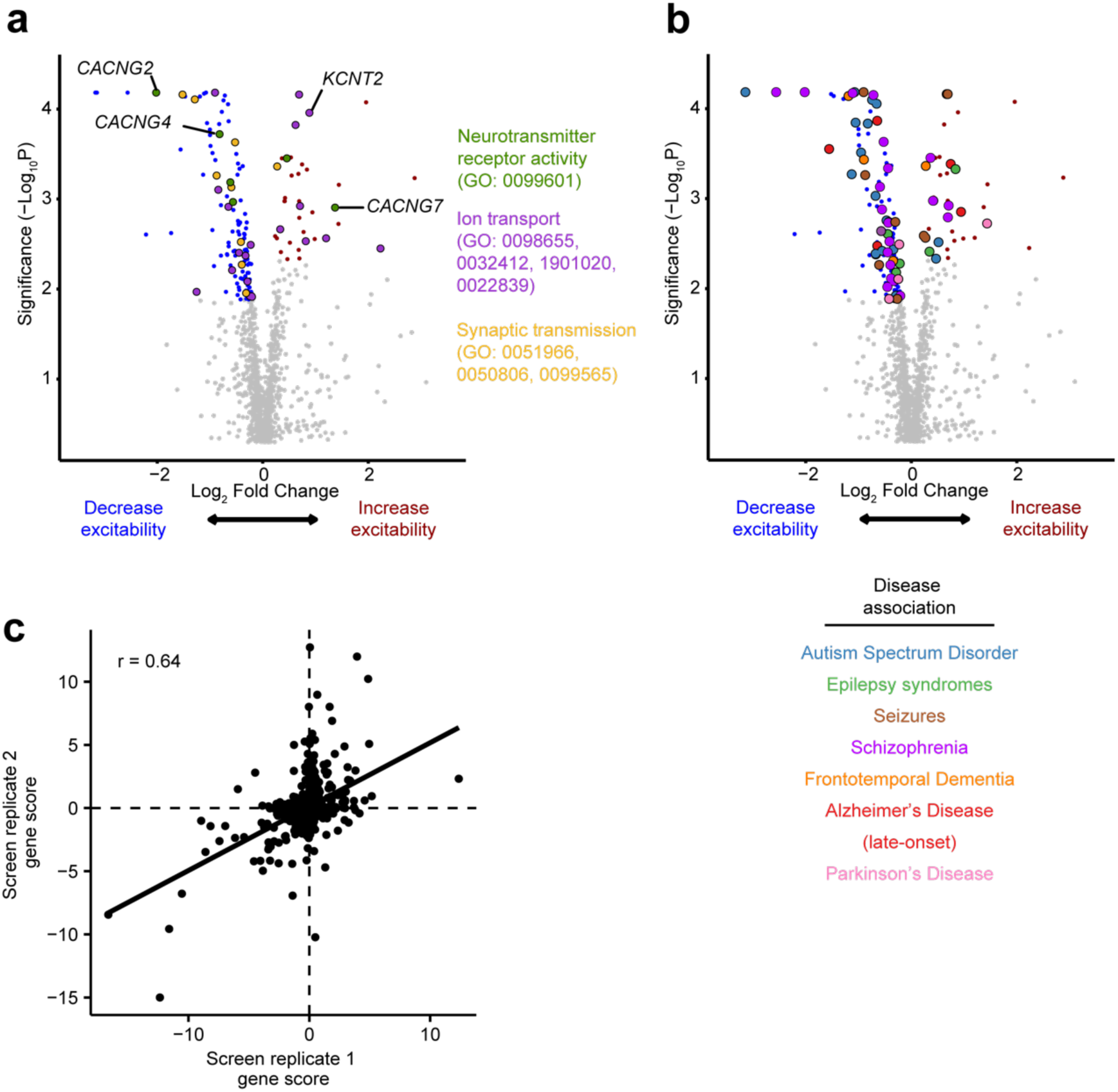
Results from Primary screen. (a) Volcano plot summarizing the effect of knockdowns on CaMPARI phenotypes and the determine statistical significance for targeted genes (Mann-Whitney U test). Dashed lines indicate a false-discovery rate (FDR) cutoff of 0.05, based on the phenotype score for gene calculated from 5 targeting sgRNAs. Black points indicate hit genes where knockdown changes CaMPARI red/green fluorescence ratio in response to glutamate stimulation, while grey points indicate non-hits. Genes related to synaptic transmission (purple) and ion transport (orange) are highlighted based on Gene Ontology terms. (b) Disease associated genes from DisGenNet and whole-exome sequencing (Satterstrom et al)9 overlap with hit genes from CaMPARI screen. (c) Scatterplot of gene scores from each replicate of the primary screen shown the correlation between screens. Pearson’s R = 0.49, linear regression fit depicted by solid black line.

**Extended Data Figure 4.**
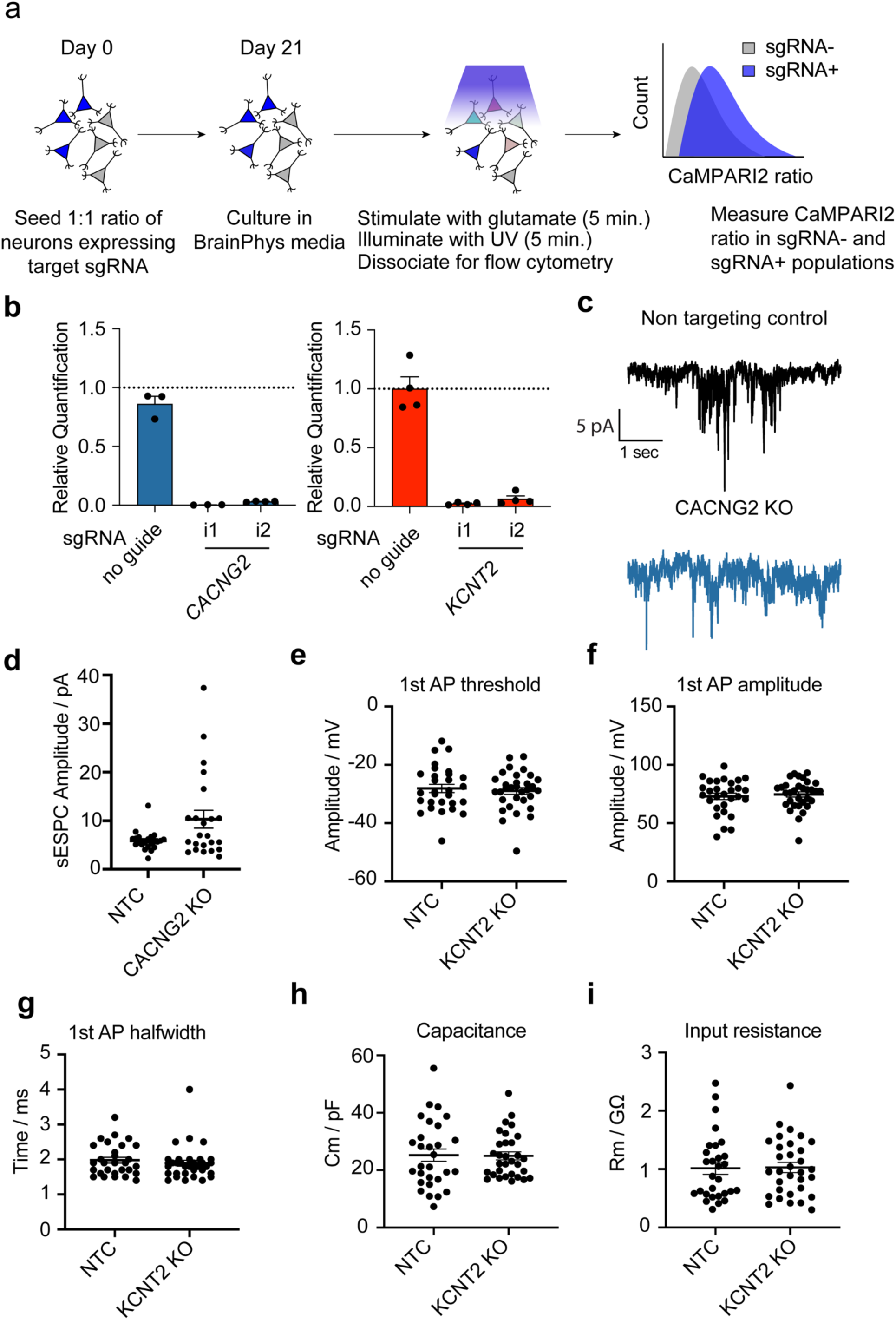
Electrophysiology profiles of iNeurons with KCNT2 and CACNG2 knockdowns. (a) Schematic of intra-well comparison experiments with 50% of CaMPARI2 iNeurons expressing a sgRNA for a target gene. A 1:1 ratio of iNeurons expressing the sgRNA: not expressing a sgRNA are seeded and differentiated. These are then treated with the same glutamate and photoconversion conditions. These are gated based on the presence of BFP in the sgRNA construct, and the CaMPARI2 ratio is directly compared between the two populations. (b) RT-qPCR shows the knock down efficiency of the sgRNAs targeting CACNG2 (blue) and KCNT2 (red). N = 4 biological replicates per condition. Error bars are SEM. (c) Sample sEPSC recordings from NTC (black trace) and CACNG2 KO (blue trace) iNeurons (d) CACNG2 KO iNeurons show a slight increase in sESPC amplitude that is not statistically significant. (NTC: 22 neurons, CACNG2 KO:23 neurons, Mann-Whitney test. (e-i) No differences between NTCs (N= 29 neurons) and KNCT2 KOs (N = 32 neurons) in either passive membrane properties or action potential kinetics. Error bars donate standard error.

**Extended Data Figure 5.**
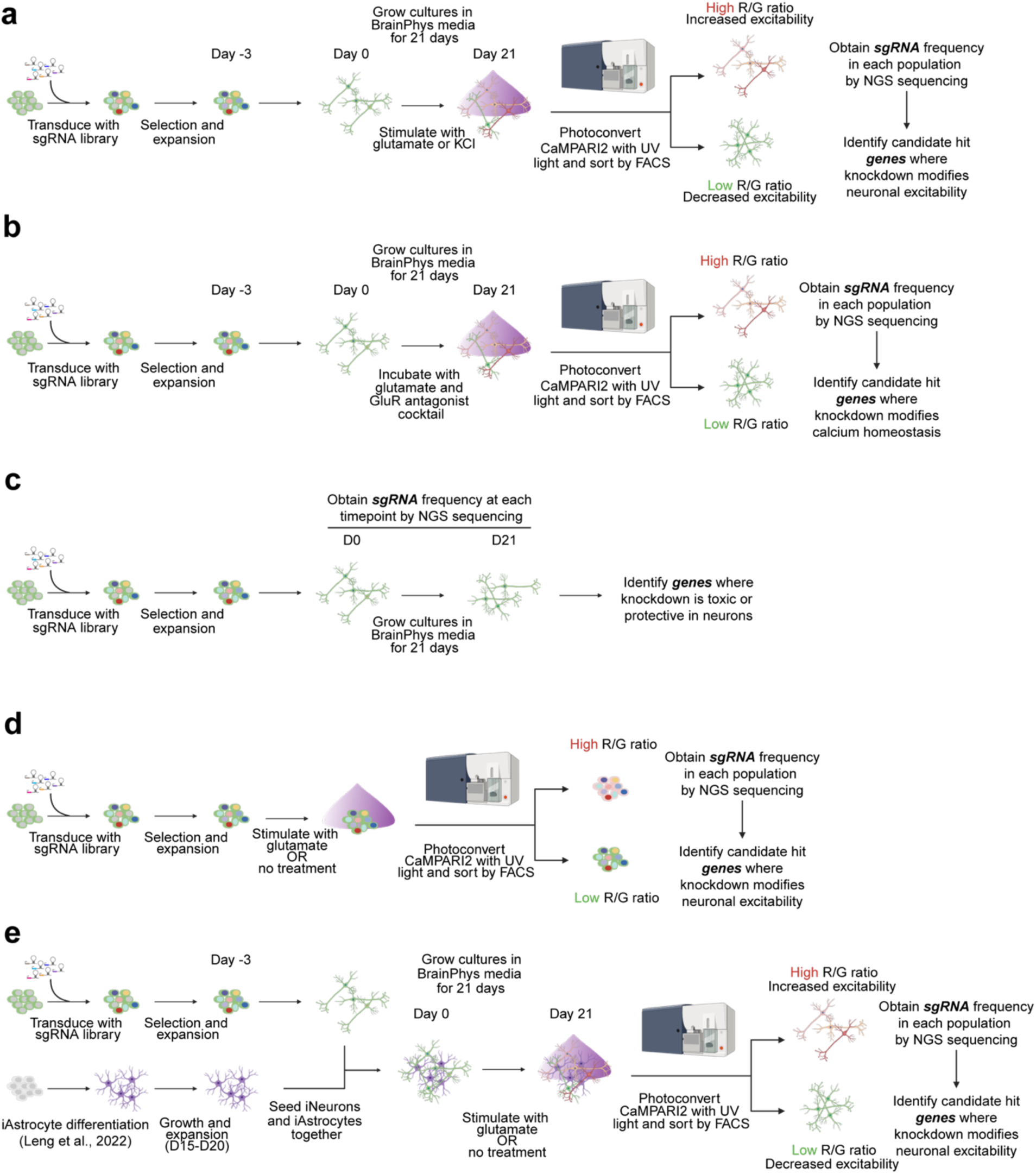
Secondary CRISPRi screen strategies. (a) Screening strategy. CRISPRi iPSCs were transduced with a pooled lentivirus sgRNA library consisting of hit genes from the primary screen (1622 sgRNAs targeting 424 genes). Exposure to puromycin selects for iPSCs successfully transduced with sgRNA construct. iPSCs were then differentiated into neurons via overexpression of NGN2 and cultured for 21 days. Neurons were then incubated with either 30 µM glutamate or 50 mM KCl, illuminated with UV light, and then separated by FACS into populations with high versus low red/green fluorescence ratios. (b) iPSCs were differentiated into neurons via overexpression of NGN2 and cultured for 21 days. Neurons were then incubated with 30 µM glutamate, 1.5 µM TTX, 10 µM NBQX, 50 µM APV, 100 µM LY367385, 0.1 µM LY341495, 1 µM CPPG, 1 µM MTEP. Neurons were then illuminated with UV light, and then separated by FACS into populations with high versus low red/green fluorescence ratios. (c) Survival of neurons was assessed by comparing sgRNA frequency in neurons at the start of differentiation (day 0) and after 21 days of differentiation. (d) iPSCs expressing sgRNAs were treated with 30 µM glutamate for 5 minutes, illuminated with UV light, and then separated by FACS into populations with high versus low red/green fluorescence ratios. (e) iPSCs expressing sgRNAs were differentiated into neurons and combined with Day 20 iAstrocytes. These co-cultures were grown for 21 days. Neurons were then treated 30 µM glutamate, illuminated with UV light, and then separated by FACS into populations with high versus low red/green fluorescence ratios. To assess the spontaneous activity of these cultures, no glutamate stimulation was applied.

**Extended Data Figure 6.**
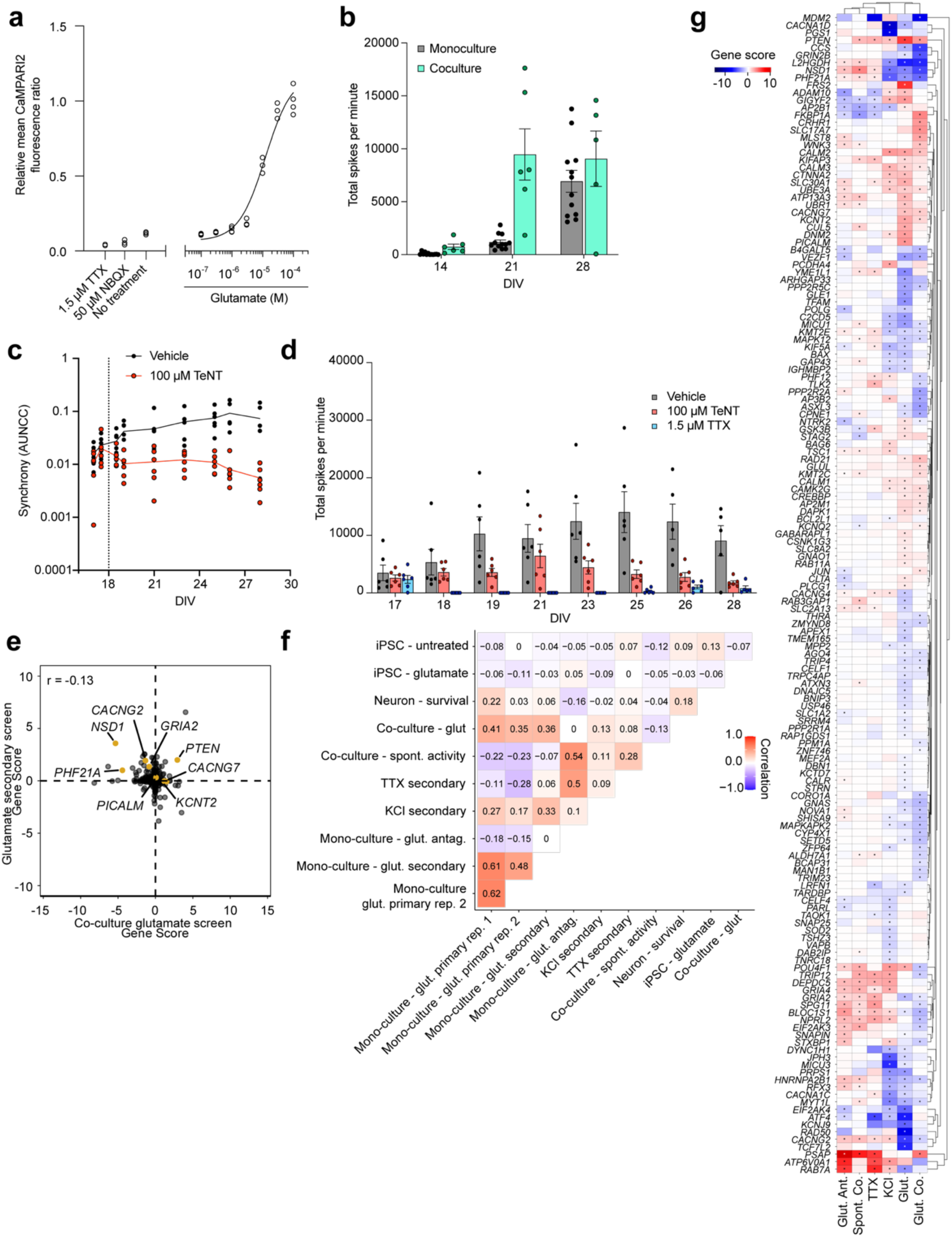
Secondary CRISPRi screens reveal excitability phenotypes in iPSC neurons. (a) CaMPARI2 neurons in coculture with iAstrocytes respond to glutamate in a dose-dependent manner. N = 4 independent culture wells per measurement. TTX (1.5 µM), NBQX (50 µM), and “no treatment” conditions did not contain glutamate. (b) Multielectrode arrays (MEA) recordings of spikes per minute of neuron-astrocyte cocultures and monoculture neurons. After 21 days of culture, more spikes are detected in cocultures. N = 6 independent culture wells, each with 16 electrodes. (c) Synchrony of neuronal activity in neuron-astrocyte cocultures on MEA as measured through Area Under a Normalized Cross Correlation. After recording on day 18 (dotted line), cultures were treated with 100 µM tetanus toxin (TeNT) or vehicle control. Loss of synchrony begins around 24 hours later in TeNT treated wells, while vehicle treated wells show a slight increase. N = 6 independent culture wells, with 16 electrodes per well. (d) Total spikes per minute of cultures from (c). Day 18 recording measured before addition of TeNT. N = 6 independent culture wells, with 16 electrodes per well. (e) Modifiers of activity of spontaneously firing neurons in coculture do not correlate with those of glutamate-stimulated neurons in coculture (Pearson’s R = –0.13) (f) Correlation matrix of gene scores from CaMPARI2 screens, using Pearson’s correlation. (g) Heatmap of gene scores in secondary screens. Each included gene is classified as a hit in at either the KCl or glutamate (monoculture and coculture) screens (FDR > 0.1) and is denoted with an asterisk when classified as a hit.

**Extended Data Figure 7.**
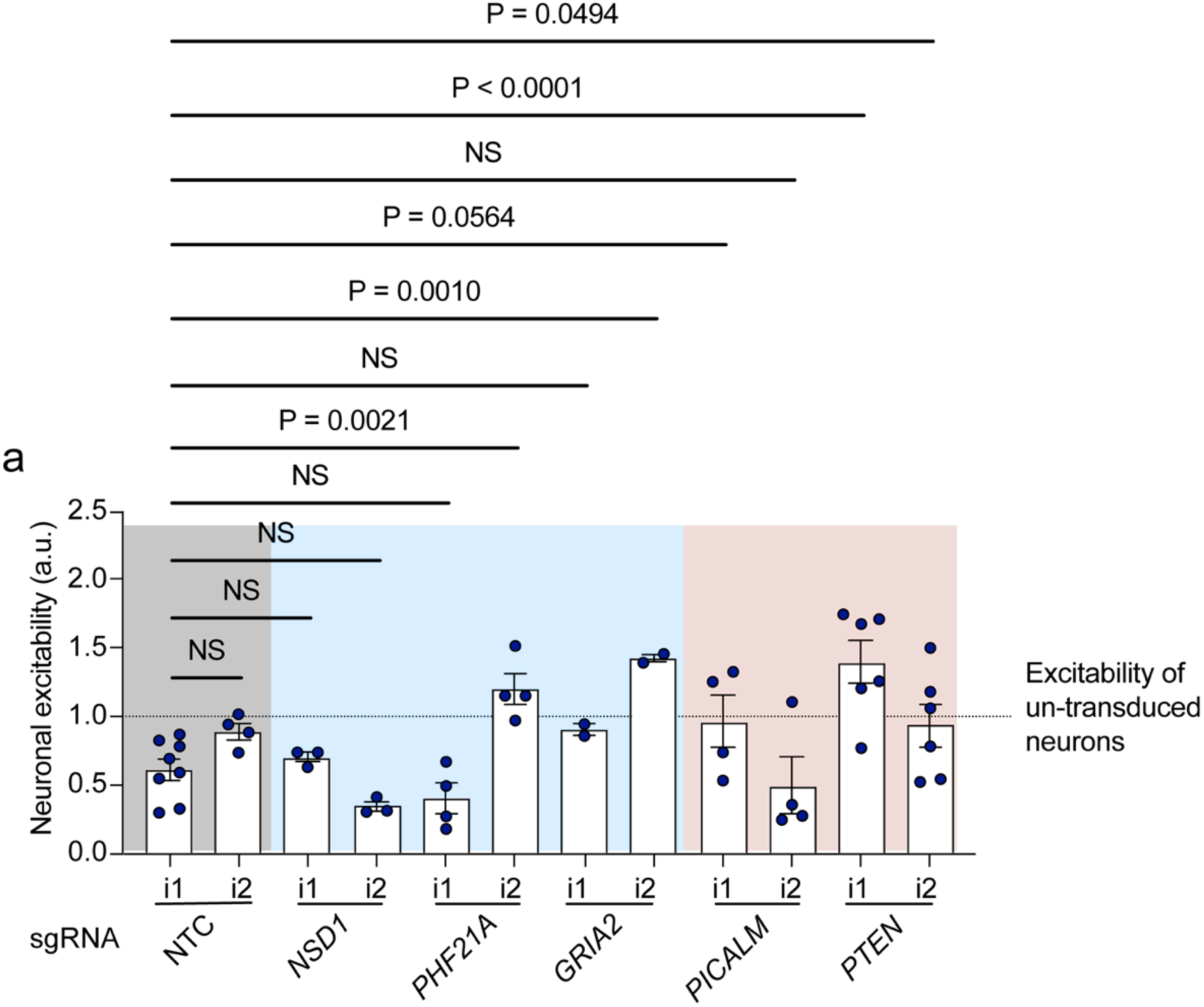
Validation of KDs from secondary screens. In cultures where 100% of CaMPARI2 iNeurons are expressing sgRNAs targeting NSD1 and PHF21A, these iNeurons lack statistically significant reduced neuronal excitability in response to glutamate stimulation observed in the screen relative to non-targeting controls. iNeurons expressing sgRNAs targeting PTEN show the expected increase in activity. For iNeurons expressing sgRNAs targeting GRIA2 and PICALM, the expected screen phenotype is not preserved when 100% of the cells have the knockdown. CaMPARI2 ratios are normalized to neuron samples that do not receive a sgRNA and are run on the flow cytometer on the same day. N = 2-8 independent culture wells per experiment. Each individual experiment was normalized to the excitability (CaMPARI2 response) of wells where neurons did not receive a sgRNA. Error bars denote standard error. P values calculated by ANOVA, using NTC_i1 as reference. P > 0.05 = not significant (NS).

**Extended Data Figure 8.**
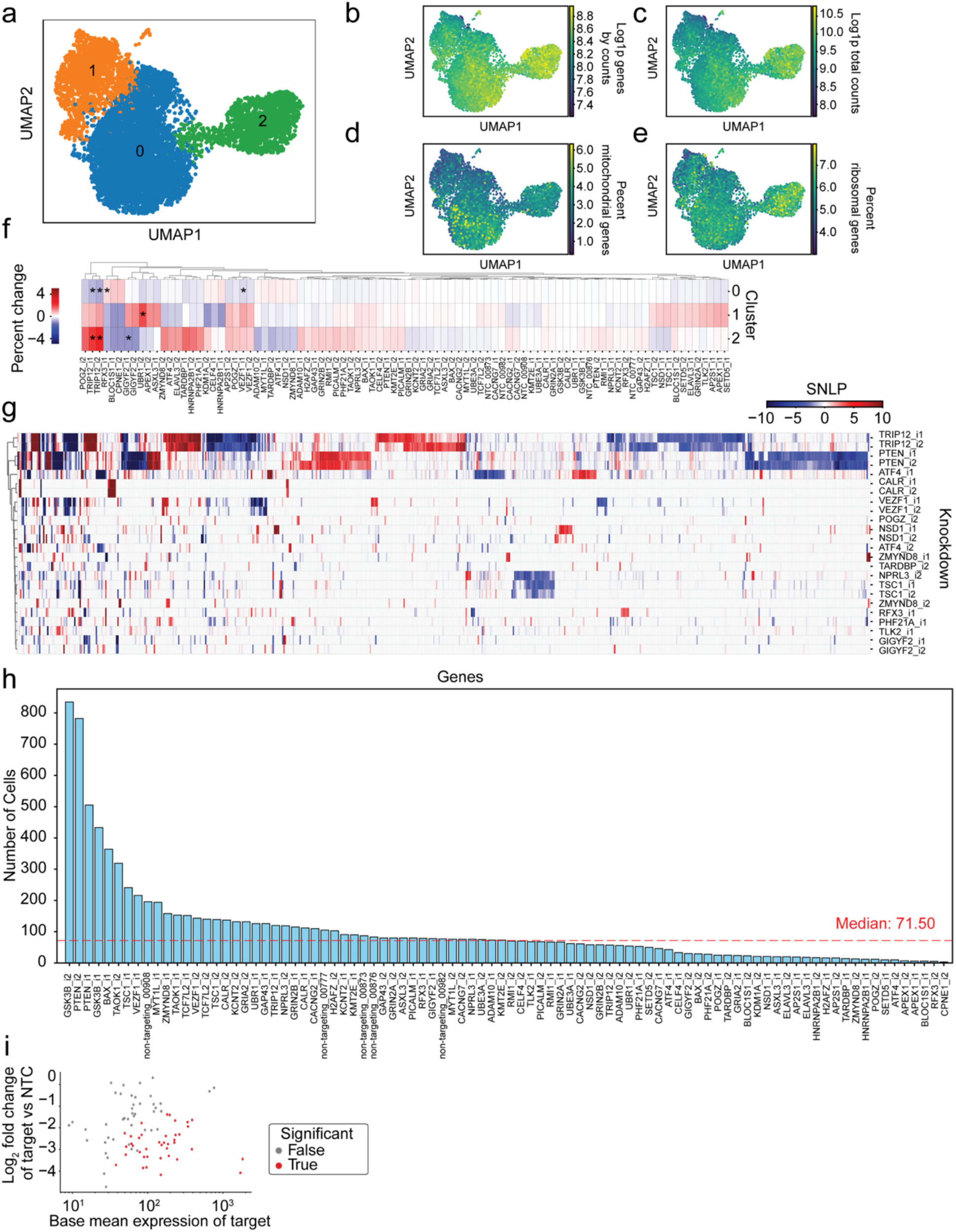
CROP-seq reveals transcriptional changes to genes involved in synaptic processes drive changes in excitability. (a) Leiden clustering of neurons from CROP-seq classified into 3 clusters. (b) Log(1+P) genes detected in each cell. (c) Log(1+P) total read counts in each cell. (d) Percentage of mitochondrial genes detected in each cell. (e) Percentage of ribosomal genes detected in each cell. (f) Heatmap of cluster shift analysis for each knockdown. Color values are expressed as the percent change in abundance of the knockdown in each cluster relative to non-targeting controls. Asterisks denote significant enrichments and depletions (adjusted q-value < 0.05). (g) Heatmap of differentially expressed genes (DEGs, versus non-targeting controls, adjusted p-value < 0.05) of knockdowns with significant on-target KD. Clustering shows consistency between individual sgRNAs targeting the same gene. SNLP is signed negative log of the adjusted p-value. Knockdowns with >10 DEGs are shown. (h) sgRNA representation by number of neurons detected in scRNA-seq data post quality control filtering and with a sgRNA MOI of one. (i) Plot of on-target KD versus base mean expression for each sgRNA. Lower expression of genes generally resulted less ability to call significance.

