## Supplementary Fig. 1 for "A Massively Parallel CRISPR-Based Screening Platform for Modifiers of Neuronal Activity"

**Supplementary Information**

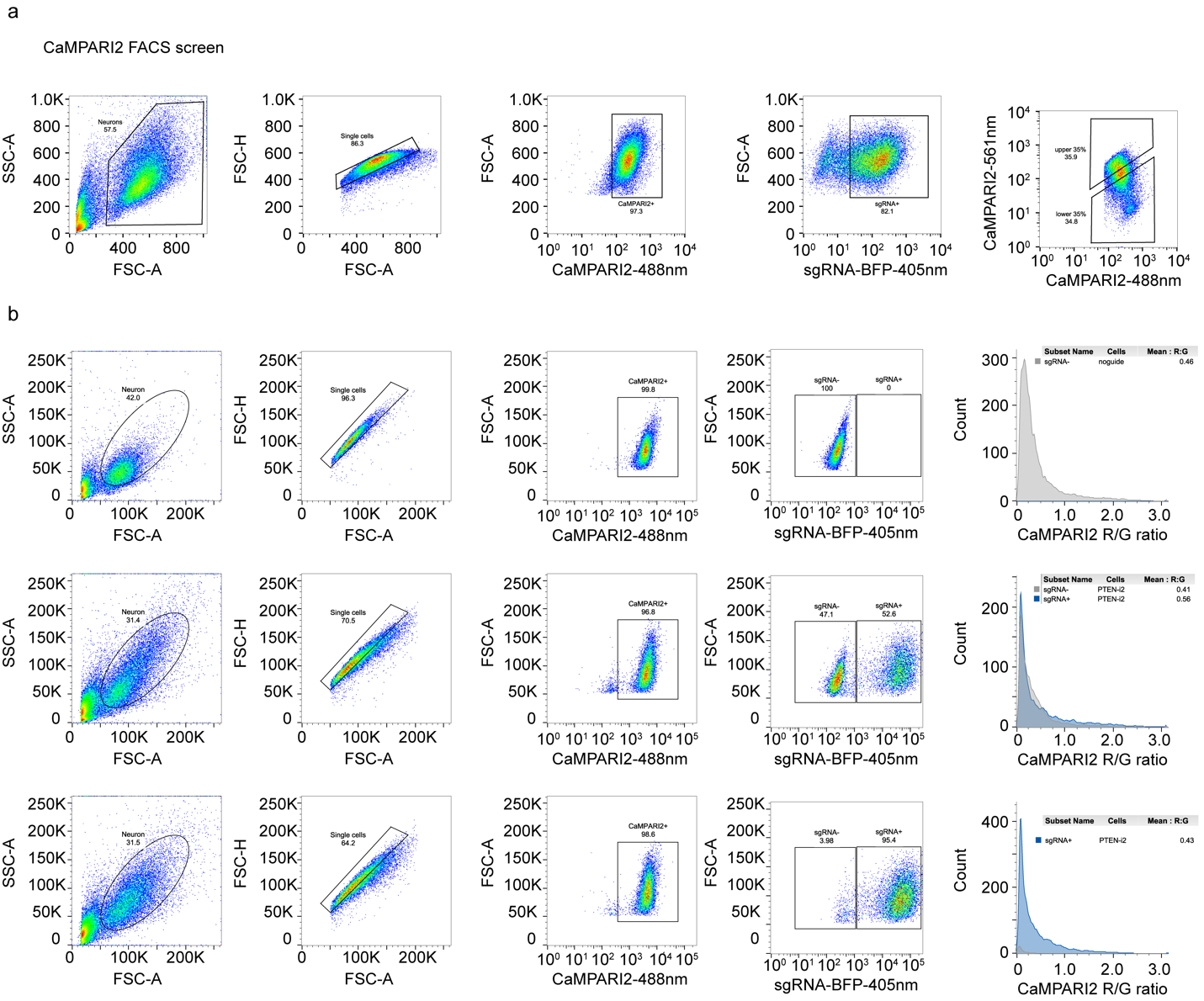

**Supplementary Figure 1. Gating strategies for FACS-based screens and for CaMPARI2 ratio validation experiments.**

a,b. Gating strategies for (a) CaMPARI2 screen and (b) CaMPARI2 flow cytometry analysis of neuronal excitability. Intact iPSC-neurons were identified from FSC-SSC plot and then gated for singlets. These cells were then gated for CaMPARI2 and BFP (sgRNA) expression. For FACS screen (a) these cells were then sorted into high and low CaMPARI2 ratio populations corresponding with the top 35% and bottom 35% of the signal distribution. (b) For flow analysis, the mean CaMPARI 2 R/G ratio was measured for BFP- (sgRNA-) and BFP+ (sgRNA+) cells.
